# Distributed fMRI patterns coupled to low-frequency cardiorespiratory dynamics provide markers of aging

**DOI:** 10.64898/2025.12.15.694468

**Authors:** Shiyu Wang, Richard Song, Laurent M. Lochard, Jiawen Fan, Yamin Li, Kimberly Rogge-Obando, Caroline Martin, Sarah E. Goodale, Haatef Pourmotabbed, J. Mason Harding, Terra Lee, Chang Li, Shengchao Zhang, Roza G. Bayrak, Taylor Bolt, Jason S. Nomi, Lucina Q. Uddin, Jingyuan E. Chen, Mara Mather, Catie Chang

## Abstract

How aging affects brain-body connections can be investigated through changes in the coupling between functional magnetic resonance imaging (fMRI) signals and bodily autonomic processes across the adult lifespan. Recent studies using univariate approaches have identified age-related changes in the association between fMRI signals from multiple individual brain regions and low-frequency respiratory and cardiac activity. Here, we investigate if whole-brain spatial fMRI patterns associated with low-frequency physiological processes (heart rate and respiratory volume fluctuations) present generalizable changes with age. Data from human participants of both sexes are included in the analysis. We find that chronological age can be predicted statistically beyond chance from patterns of low-frequency fMRI-physiology coupling, even after accounting for individual differences in physiological signal characteristics and brain anatomy. Notably, brain areas implicated in central autonomic regulation, including nodes within salience and ventral attention networks (e.g., insula and middle cingulate cortex), are amongst the strongest contributors to age prediction. Further, we observe that after removing physiological effects from fMRI data, the residual blood oxygen level-dependent (BOLD) signal variability is still a reliable indicator of age. Together, these findings underscore the close integration between brain and body physiology, and highlight this interaction as a potential biomarker of the aging process.

**Significance Statement:** The association between brain activity and respiratory or cardiac activity is often dismissed as “noise” in functional magnetic resonance imaging (fMRI) studies. However, emerging evidence suggests that coupling between fMRI and peripheral physiological signals can provide valuable insight into the brain-body connection. In this study, we show that whole brain patterns of coupling between fMRI signals and low-frequency respiratory and cardiac processes can reliably predict age across the adult lifespan. Brain regions involved in autonomic regulation, such as insula and cingulate cortex, were among the most informative predictors of age. These findings suggest that fMRI-physiology coupling may capture aging-related changes in brain vascular health and autonomic function and may have broader relevance for tracking disease-related disruptions in brain–body interaction.

## Introduction

Physiological processes in the body have long been known to influence functional magnetic resonance imaging (fMRI) signals. For example, breathing and cardiac activity introduce fMRI artifacts time-locked to each breath and heartbeat that are unrelated to blood oxygen changes (Glover et al., 2000; Murphy et al., 2013). However, beyond these cyclic artifacts, slow variations in breathing and cardiac activity are associated with low-frequency fluctuations in the blood oxygen level-dependent (BOLD) fMRI signal. Mechanisms linking systemic low-frequency physiological signals to BOLD signals include neuronal autonomic activity (Tu and Zhang, 2022), changes in cerebral blood flow driven by fluctuating arterial CO_2_ concentrations (Wise et al., 2004; Birn et al., 2006), and sympathetically mediated vasoconstriction (Duyn et al., 2020; Picchioni et al., 2022). Though long considered “noise” (Uddin, 2020), low-frequency BOLD fluctuations coupled with slow variations in peripheral physiology exhibit network-like structures (Bright et al., 2020; Chen et al., 2020) and robust global spatiotemporal responses (Bolt et al., 2025). Increasing evidence further highlights the role of cardiorespiratory activity in regulating neuronal activity (Zelano et al. 2016; Goheen et al. 2023).

Importantly, the relationship between systemic physiology and fMRI signals may change with aging, potentially reflecting age-related alterations in brain vasculature or autonomic function (e.g., (Mayhan et al., 1990; Riecker et al., 2003; Handwerker et al., 2007; Ungvari et al., 2008; Kannurpatti et al., 2010; Thayer et al., 2010; Flück et al., 2014; Thomas et al., 2014; McKetton et al., 2018; Chen, 2019; Kumral et al., 2019; Yabluchanskiy et al., 2021; Delli Pizzi et al., 2023; Giunta et al., 2024; Mather, 2024). Older adults exhibit smaller BOLD responses to breath-holding tasks (Handwerker et al., 2007) and altered relationships between resting heart rate variability (HRV) and functional connectivity of autonomic control regions (Kumral et al., 2019). These studies suggest that BOLD components coupled to systemic physiology may carry useful information regarding the aging brain.

In a separate line of work, aging has been linked with changes in overall BOLD signal variability, a measure hypothesized to reflect neural efficiency and flexibility (Yan et al., 2011; Garrett et al., 2013b; Grady and Garrett, 2014; Guitart-Masip et al., 2016; Nomi et al., 2017; Waschke et al., 2021; Zhong and Chen, 2022). Regional BOLD variability is reported to exhibit linear and quadratic relationships with age (Garrett et al., 2010; Nomi et al., 2017; Goodman et al., 2024), and multivariate BOLD variability patterns may be predictive of age (Millar et al., 2020). However, whether the specific component of fMRI variability coupled to low-frequency physiological fluctuations is itself predictive of age is unclear. Recent studies using univariate, correlation-based approaches have begun to identify age-related differences in how low-frequency physiology relates to fMRI (Fan et al., 2025; Song et al., 2025); but none have tested if brain-wide multivariate patterns of low-frequency physiology-fMRI coupling can reliably predict age. Such predictability would suggest that distributed age-related changes in fMRI-physiological coupling have generalizable signatures.

Here, we aim to determine whether large-scale patterns of fMRI coupled to low-frequency physiological processes (“physio-fMRI coupling”) contain reliable signatures of aging. We use a multivariate modeling framework to predict chronological age from the portion of fMRI variability coupled to low-frequency physiology, and to characterize how these brain-body interactions evolve with age. We design permutation tests to establish that observed effects are not driven solely by age-related changes in respiratory or cardiac activity, low-frequency fMRI spectral content or brain anatomy. We then investigate nonlinearities in the relationship between physio-fMRI coupling and age, motivated by studies showing nonlinear age effects in the spatial topology of global signal (Nomi et al., 2024) and in the respiration- or cardiac-related temporal lags with cortical global signal (Fan et al., 2025). Finally, we examine whether previously reported age effects on overall BOLD variability can be attributed to low-frequency physiological effects. This work has implications for novel biomarkers of aging and highlights the biological significance of the physiological “noise” (Uddin, 2020) in fMRI.

## Materials and Methods

### Datasets

Data used in this study were drawn from three publicly available databases with concurrent fMRI and physiological recordings: (1) the Nathan Kline Institute-Rockland Sample (NKI) dataset (Nooner et al., 2012); (2) the Human Connectome Project Lifespan Sample (HCPA) dataset (Harms et al., 2018); and (3) the Heart Rate Variability - Emotion Regulation (HRV-ER) dataset (Yoo et al., 2023). All participants included in the study are cognitively unimpaired and are considered healthy adults. The participant characteristics, acquisition, and pre-processing steps for each dataset are described below.

#### fMRI acquisition and preprocessing: NKI dataset

The 3T resting-state fMRI data from the Nathan Kline Institute-Rockland Sample (NKI) dataset were acquired with voxel size = 2 mm isotropic, TR = 1400 ms, 404 volumes (scan duration ≈ 9.4 min) (Nooner et al., 2012). We included 161 subjects (19-83 years old, 102 females) with resting-state fMRI data and concurrently recorded respiratory and cardiac signals. The preprocessing of the resting-state fMRI data included motion correction (FSL *mcflirt*) and ICA-based FIX-correction to control for motion and other non-BOLD noise (Salimi-Khorshidi et al., 2014). For the latter, we selected 25 subjects randomly in an age-balanced manner, and hand-labeled the melodic ICA components to create the NKI specific FIX ICA training set. For certain analyses, the data were also registered to MNI152 template space using ANTS (Avants et al., 2011), followed by spatial smoothing with a Gaussian filter of 3 mm full-width-half-maximum (FWHM).

#### fMRI acquisition and preprocessing: HCPA dataset

The 3T HCPA resting-state fMRI dataset was acquired with the following parameters: voxel size = 2 mm isotropic, TR = 800 ms, 487 volumes, scan duration ≈ 6.5 min (Harms et al., 2018; Bookheimer et al., 2019). In total, 1805 scan sessions from 577 subjects (age range 36-89 years old, 323 females) were included in the study that passed our physiological signal quality assessment (described below). The resting-state fMRI that we used had already gone through the HCP generic fMRI volume preprocessing pipeline, was denoised using ICA-based FIX-correction, and was registered into the MNI152 template space (Glasser et al., 2013; Salimi-Khorshidi et al., 2014).

#### fMRI acquisition and preprocessing: HRV-ER dataset

The resting-state fMRI scans from the HRV-ER dataset were acquired using a multi-echo echo-planar imaging sequence with the following parameters: TR = 2400 ms, TE 18/35/53 ms, voxel size = 3.0 mm isotropic, 175 volumes, scan duration = 7 min (Yoo et al., 2023). We included the pre-intervention scan sessions from 59 subjects from the younger adult group (18-30 years old, 35 females) and 51 subjects from the older adult group (55-80 years old, 35 females). The preprocessing for the HRV-ER dataset included motion correction, slice timing correction, denoising with the multi-echo independent component analysis (ME-ICA) based method *tedana* to remove non-BOLD components, spatial registration to MNI152 template space, and smoothing with 3 mm FWHM (Kundu et al., 2012, 2017).

#### fMRI ROI extraction: NKI, HCPA, and HRV-ER datasets

The mean fMRI time courses from the following ROIs were extracted: 400 cortical ROIs from the Schaefer Atlas (17 networks) (Schaefer et al., 2018), 16 subcortical ROIs from the Melbourne Atlas (3T scale) (Tian et al., 2020), 9 brainstem ROIs from the Harvard Ascending Arousal Network Atlas version 1.0 (Edlow et al., 2012), and 72 white matter ROIs from the Pandora Tractseg Atlas with a 0.95 threshold (HCP version) (Hansen et al., 2021). Features extracted from white matter functional data have been found to change with aging (Li et al., 2023, 2024; Xu et al., 2024). In addition, white matter BOLD signal has a close association with systemic physiological variations (Özbay et al., 2018), and adding white matter ROIs improves the reconstruction accuracy of low-frequency RV and HR signals from fMRI data (Bayrak et al., 2025). Taken together, we suspect that the coupling between white matter BOLD and low-frequency physiological processes may be related to some of the aging effects reported in recent literature, thus including white matter ROIs in our study. After obtaining the ROI time courses from each atlas, we separated the entire time course from a given scan into two overlapping 5-minute sections (TR1 - TR214 and TR191 - TR 404 for NKI, TR1 - TR375 and TR104 - TR478 for HCPA, TR1 - TR125 and TR51 - TR175 for HRV-ER), and preserved the sections with a mean framewise displacement (mean FD) smaller than 0.50 mm to control for excessive motion. On each of the retained 5-min fMRI scan segments, we then detrended the fMRI time series up to a 4th degree polynomial to control for slow scanner hardware drifts, regressed out 6 motion realignment parameters and their first-order temporal derivatives (12 motion regressors in total), excluded the first and last 5 seconds (3 frames for NKI, 6 frames for HCPA, and 2 frames for HRV-ER) to avoid potential edge effects at the boundaries of the segments, and z-normalized each ROI’s time course.

#### fMRI ROI extraction: subject-specific approach

Since changes in brain morphology that occur in aging may cause age-related errors in the alignment to a single standard template, we also examined an individualized native space parcellation approach (“IND”) using the NKI dataset. Freesurfer *recon-all* and *mri_aparc2aseg* were applied to obtain 400 subject-specific cortical ROIs from the Schaefer Atlas (Schaefer et al. 2018). The recon-all program preprocesses the subject T1-weighted anatomical image using steps including intensity normalization, skull stripping, white- and gray-matter classification, and cortical surface reconstruction, and *mri_aparc2aseg* maps the Schaefer parcellation onto the subject’s cortical surface by matching the cortical folding geometry and converting it to a native-space volumetric segmentation (Dale et al., 1999; Fischl et al., 1999, 2002).

#### Physiological signal acquisition and processing

For all datasets, peripheral physiology was monitored continuously during fMRI with a respiratory belt placed around the subject’s abdomen and a photoplethysmogram (PPG) placed on the subject’s finger (Nooner et al., 2012; Harms et al., 2018; Bookheimer et al., 2019; Yoo et al., 2023). The sampling rate of the physiological recordings is 62.5 Hz for the NKI dataset, 400 Hz for the HCPA dataset, and 10 kHz for the HRV-ER dataset. All physiological signal quality assessment was done manually. For the HCPA dataset, the quality assessment was cross-checked with 2 researchers. For the NKI dataset, two rounds of physiological signal quality assessment were done with 7 researchers. For the HRV-ER dataset, only PPG data was considered for further analysis, since the respiratory belt recording was corrupted for many of the scans. End-tidal CO_2_ recordings are available for the HRV-ER dataset, and end-tidal CO_2_ is related but not equivalent to respiratory belt recordings. However, the HRV-ER dataset is used here only as an external test set for the age prediction models derived from NKI and HCPA datasets, and it may be difficult to directly translate models trained on features extracted from respiratory belt metrics to end-tidal CO_2_ metrics.

As shown in **Fig. 1a**, the respiratory belt and PPG signals were preprocessed as follows: 1) the physiological recordings were aligned with the fMRI triggers, 2) respiratory variation (RV) was calculated as the standard deviation (SD) of a sliding window of 6 s centered at each fMRI TR, and heart rate (HR) was calculated from the PPG signal as the inverse of the mean inter-beat-interval in the same window (Chang et al., 2009; Chen et al., 2020), and 3) both RV and HR were divided into 5-minute segments aligning with those of the fMRI ROIs, and were then detrended up to a 4th degree polynomial. The first 5 and last 5 seconds of each segment were excluded, and the resulting segments were z-normalized. Heart rate variability measures for each 5-minute segment, calculated as the root mean square of successive differences (rMSSD, capturing higher-frequency HRV), was calculated before detrending.

**Figure 1.**
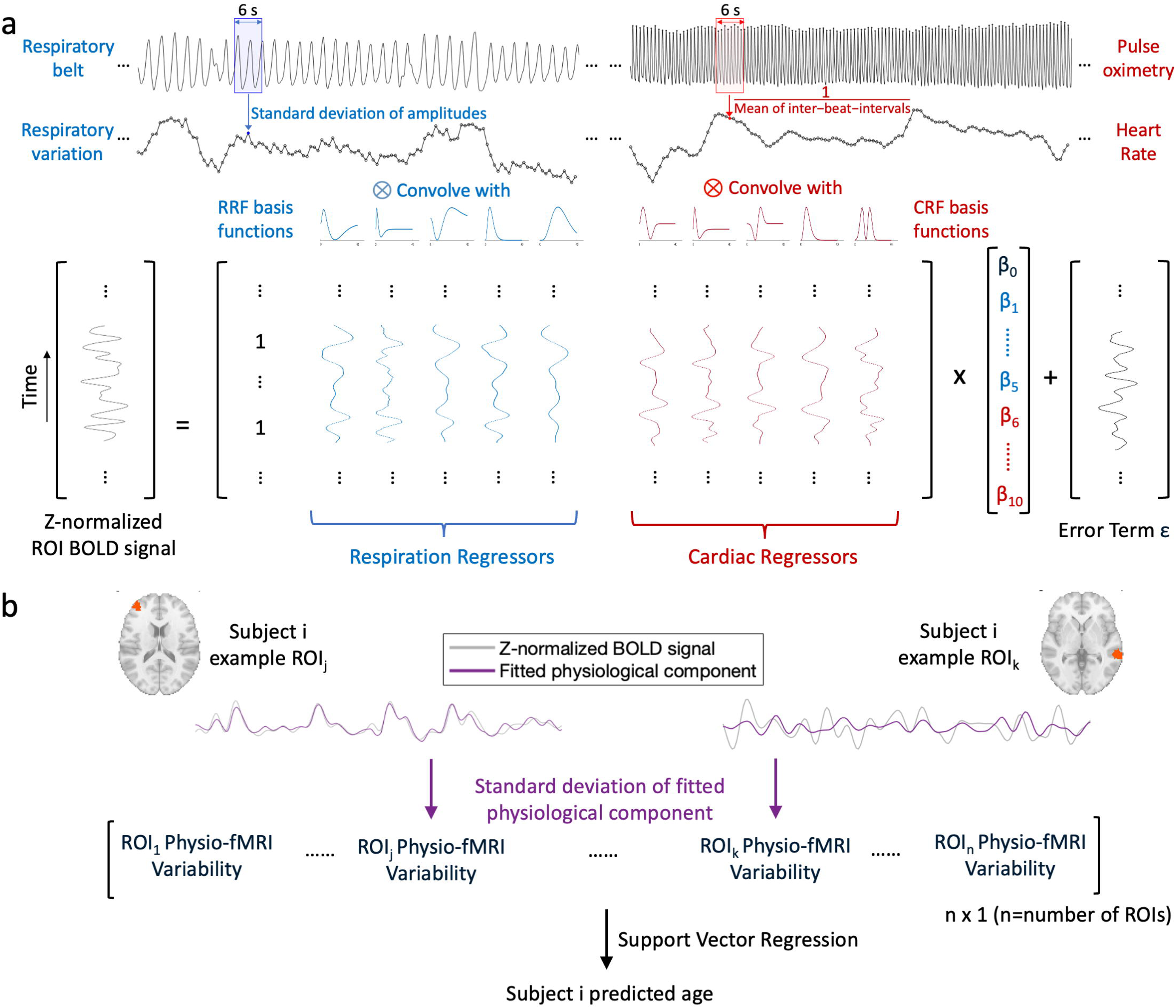
Flowchart showing the framework for predicting age from fMRI physiological variability. a) Respiratory variation (RV) and heart rate (HR) time courses are extracted from the respiratory belt and pulse oximetry recordings, respectively, using 6-s sliding windows centered at each fMRI TR. RV and HR signals are then detrended up to a 4th order polynomial (not shown in the figure) and convolved with their corresponding response function basis sets (5 respiratory response basis functions (RRF), and 5 cardiac response basis functions (CRF)). The convolved signals are fitted to the fMRI BOLD signal. b) Standard deviation of the fitted physiological component, here referred to as the physio-fMRI variability, is then computed for each ROI. This measure is equivalent to the square root of percent variance explained in the BOLD signal by the physiological component. The resulting collection of physio-fMRI variability values, across all ROIs, comprises the features used for age prediction via support vector regression. In our experiments, the physiological component is based on either RV only, HR only, or RV and HR jointly.

### Deriving the fMRI physiological component

The relationships between BOLD signal and low-frequency physiological signals have been modeled in healthy young adults via physiological response functions (PRFs) (Birn et al., 2006; Chang et al., 2009; Kassinopoulos and Mitsis, 2019; Chen et al., 2020). Recently, Chen et al. proposed a set of respiratory and cardiac basis functions that allowed flexible fit to the resting-state fMRI physiological component (Chen et al., 2020). Here we adopted the method proposed by Chen et al., convolving RV and HR each with their basis functions and fitting the resulting convolved signals to each ROI’s time course either separately (to extract the RV- or HR-specific BOLD components, respectively), or together (to obtain the RVHR jointly fitted BOLD component) (Chen et al., 2020) (**Fig. 1**). One important caveat was that these physiological response functions were developed with Gaussian temporal smoothness priors (Chang et al., 2009; Chen et al., 2020), which means that by convolving these functions with physiological signals in the time domain, only the low-frequency components were preserved in the convolved signals. Thus, the fitted fMRI signal largely reflects the coupling between physiological processes and fMRI BOLD signals in the low-frequency range (about 90% power falls between 0.01 - 0.15 Hz).

### BOLD Variability Measures

The fMRI physiological component was first derived, as described above, by fitting the set of convolved RV and/or HR signals to the temporally z-scored fMRI signal from each ROI. The standard deviation (SD) (Garrett et al., 2010, 2013a) of this fitted signal was then computed to measure its temporal variability (**Fig. 1b**). We refer to the SD of the fitted low-frequency RV, HR, or joint (RV, HR) signals as RV-fMRI variability, HR-fMRI variability, and RVHR-fMRI variability, respectively. These ROI-based variability features were then used as the input for age prediction, as described in the next section. We note that the SD of the fitted physiological signal corresponds to the (square root of the) percent variance explained in an fMRI signal by the physiological data, as the initial fMRI overall signal variance has been normalized to 1. While rMSSD is another widely used metric to quantify fMRI variability (Garrett et al., 2013b; Nomi et al., 2017; Boylan et al., 2021; Goodman et al., 2024), we chose not to apply it here since 1) the fitted fMRI physiological component is already in the 0.01-0.15 Hz low-frequency range, limiting the potential rMSSD range, 2) differences in the temporal sampling rate (TR) across datasets make it difficult to directly compare their respective rMSSD values, especially on the non-normalized data.

In addition to quantifying the portion of fMRI variability that is coupled to physiology, we also calculated measures of BOLD variability on the original preprocessed ROI time courses. On the non-z-scored fMRI data (RAW), we computed SD, ALFF (Biswal et al., 1995; Zang et al., 2007), and fALFF with a low-frequency range set to 0.01-0.08 Hz (Zou et al., 2008). For fALFF specifically, the fraction was calculated by taking the ratio of the low-frequency power to the power from 0.01-0.30 Hz (rather than up to a signal’s highest frequency) to account for the difference in the total frequency range between NKI and HCPA dataset, which have different TRs. ALFF and fALFF characterize the amplitude or proportional amplitude of fMRI BOLD low-frequency fluctuations (Biswal et al., 1995; Zang et al., 2007)(Biswal et al., 1995; Zang et al., 2007; Zou et al., 2008)(Biswal et al., 1995; Zang et al., 2007), and have been used to study the aging process (Hu et al., 2014; Yang et al., 2021; Montalà-Flaquer et al., 2022) as well as patients with cognitive impairments (Wang et al., 2021; Zhang et al., 2021), neurodevelopmental disorders (Karavallil Achuthan et al., 2023) or mental disorders (Zang et al., 2007; Egorova et al., 2017).

In addition, we regressed out the physiological components from the non-z-scored ROI time courses (REG), and computed ALFF and fALFF on the resulting signals to test if removing physiological components of fMRI data impacts the relationship between BOLD variability and aging. The SD of REG signal was not tested here since it is redundant with the RAW SD and percent variance explained by physiological components (SD^2^ = SD^2^ * (1 - SD^2^)).

### Age Prediction Experimental Design and Statistical Analysis

We used support vector regression (SVR) to perform age prediction with the fMRI variability measures. SVR has previously been used to predict age with BOLD variability (Millar et al., 2020). A 10-fold cross-validation framework was used, and the train/test/validation splits were performed at the subject level to prevent within-subject data leakage (∼81% training, ∼9% validation, and 10% testing). We compared the age prediction performance across three kernels: a linear kernel, a second-degree polynomial kernel (Poly2), and a nonlinear RBF (radial basis function) kernel. The hyperparameters C and epsilon were selected for each kernel based on the best performance (lowest mean absolute error; MAE) on the validation set. The overall age prediction performance was evaluated by first averaging the age prediction results for subjects with multiple scan segments (NKI dataset) and/or scan sessions (HCPA dataset), then concatenating the prediction results for the 10 test folds. Pearson correlation coefficient (r) and MAE between the chronological age and model-predicted age were used to quantify the prediction results. The age prediction framework was performed on the NKI and HCPA datasets. The HRV-ER dataset was used later for external validation.

For significance testing, we performed a permutation test by fitting a random physiological signal (a different scan’s RV and/or HR signal) to the fMRI signal (**Fig. 2**). In each iteration of the permutation test, we used the physiological fMRI component derived from these mismatched signals to predict either (i) the ages of subjects corresponding to the fMRI signals or (ii) the ages of subjects corresponding to the (mismatched) fitted physiological signals. This procedure was done to test that the predictive ability of the fMRI physiological components is not due to age-related changes in the physiological or fMRI time courses themselves, but rather to their coupling. We repeated the process 100 times each for the NKI native space approach, the NKI MNI-based approach, and the HCPA MNI-based approach. Significant age prediction results were determined by comparing the actual (matched) prediction MAE to the distribution of the MAE of individual permutations (100 in total) using the same model (kernel, hyperparameters) with the same train/test/validation splits (p < 0.05).

**Figure 2.**
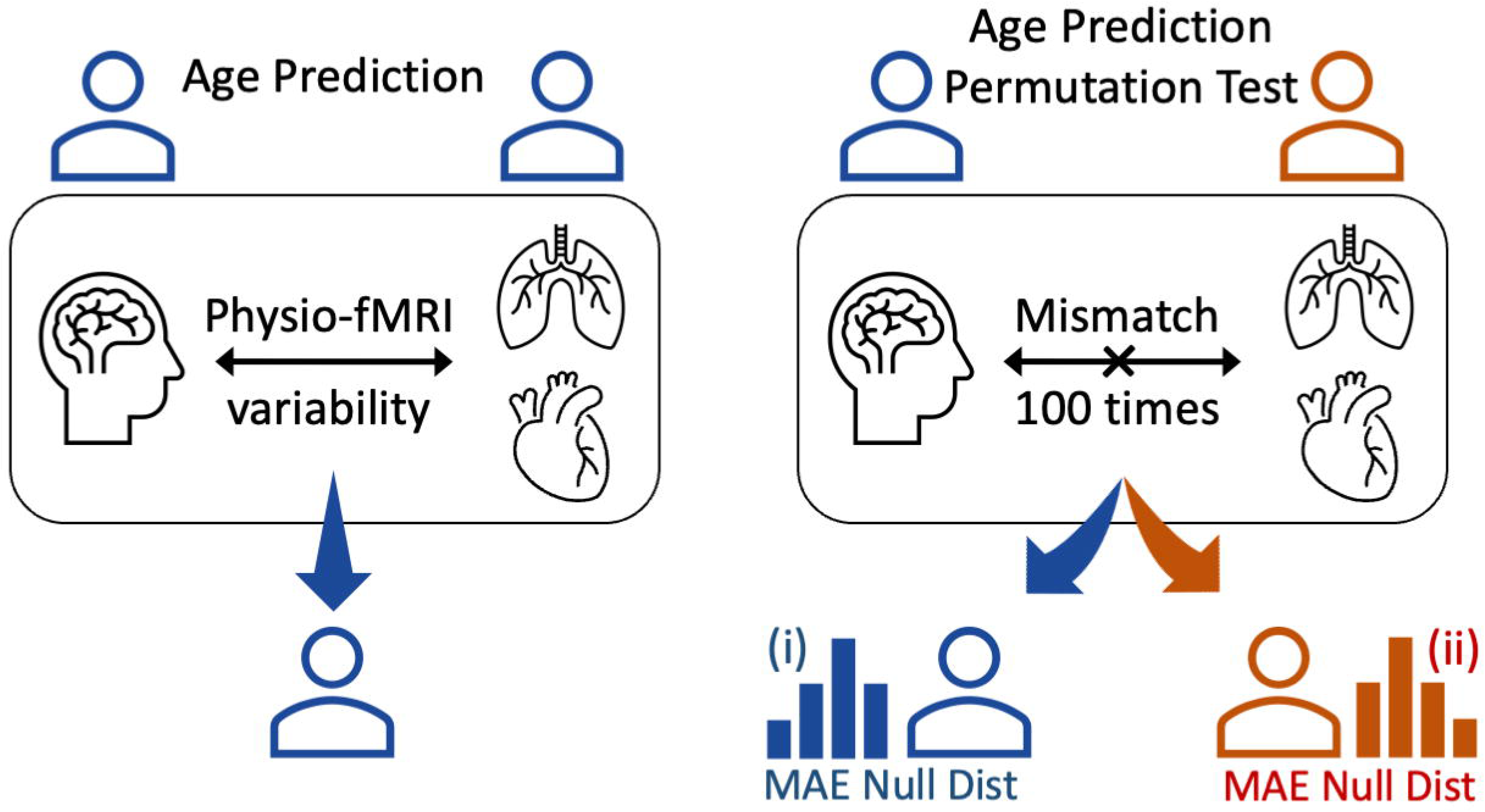
Illustration of the age prediction framework (left) and the permutation tests used to examine the significance of age prediction performance (right) with physio-fMRI variability. After pairing the fMRI signal with a mismatched physiological signal drawn from another scan (right), the same pipeline for extracting physio-fMRI variability features was applied to predict the age of the subject associated with either the (i) fMRI signal or (ii) physiological signal. This process was repeated 100 times to establish null distributions for mean absolute error (MAE Null Dist) on (i) and (ii). An age prediction result is considered to be significant if it reaches p < 0.05 with respect to both MAE null distributions, and for both the NKI and HCPA datasets.

We also compared the age prediction performance before (RAW) and after (REG) regressing out the estimated physiological component of the fMRI data. Just as above, we applied the same 10-fold cross validation procedure, with the same model and hyperparameter selection criteria, and evaluated the age prediction result by averaging the prediction results across segments corresponding to a given subject and then concatenating the 10 test folds.

Further, we compared two different training strategies using the MNI-based parcellations for the NKI and HCPA datasets: in one case, we trained two separate models for each dataset, and in the other, we jointly trained one model on two datasets combined. The exact same 10 folds were used for these two training strategies. For both strategies, performance was evaluated by concatenating the 10 test folds from the NKI and HCPA datasets.

### Assessing cortical or subcortical ROI importance for age prediction

Since it is difficult to directly interpret or visualize the nonlinear RBF kernel used for the age prediction, we used two complementary strategies to identify ROIs or groups of ROIs (networks) that were important for age prediction and to assess how they contributed to the model output. First, we used a network-based approach. The Schaefer atlas pre-assigns each of the 400 cortical ROIs to one of 17 functionally defined brain networks. For each network in turn, we compared the age prediction ability of the low-frequency fMRI physiological component using only ROIs within that network (ranging from 11-39 ROIs across networks), using the exact same 10-fold cross-validation split and model selection specified above. The second approach uses SHAP (Lundberg and Lee, 2017), a game theory based approach that approximates feature importance in complex models, which does not assume feature independence and model linearity. The SHAP value of a certain feature indicates how the model output would change when conditioning on that feature. We ran SHAP along with the 10-fold cross-validation for 400 cortical or 25 subcortical ROIs, and assessed the SHAP values of the ROI features with regard to each test sample. The SHAP analyses were performed on NKI and HCPA datasets separately as well as jointly. The SHAP values were aggregated across 10 test folds to obtain an overall summary of feature importance across the entire dataset.

### Generalizability of the age prediction framework

We directly applied models that were separately trained on all NKI subjects, or jointly trained on both datasets, to a third independent dataset (HRV-ER). Here, only the HR-coupled fMRI variance was tested since we observed that respiratory belt recordings in the HRV-ER dataset failed our quality assessment. The HRV-ER dataset contains two age groups, with 59 younger adults (18-28 years old) and 51 older adults (55-80 years old) with no middle age group in-between. Therefore, we test if the model could give significantly (p < 0.005) higher age prediction values for the older adults group compared with the younger adults, using a one-tailed unpaired *t*-test with unequal variance.

## Results

### Dataset summary statistics

The age distributions of the NKI and HCPA datasets are shown in **Figure 3a**. Heart rate variability, characterized by the root mean square of successive differences (capturing an index of parasympathetic activity), shows age-related decrease across the two aggregated datasets (**Fig. 3b**; r = -0.22, p < 0.001). Head motion during the fMRI scans, characterized by mean framewise displacement (mean FD), was weakly positively correlated with age (r = 0.24, p = 0.002 for the NKI dataset and r = 0.11, p = 0.011 for the HCPA dataset) after excluding the scan segments with excessive motion (see Methods). Subject-wise mean FD was positively correlated with physio-fMRI variability averaged across 497 cortical, subcortical, and white matter ROIs in the NKI dataset. However, the relationships were not significant in the HCPA dataset (**Table S1**).

**Figure 3.**
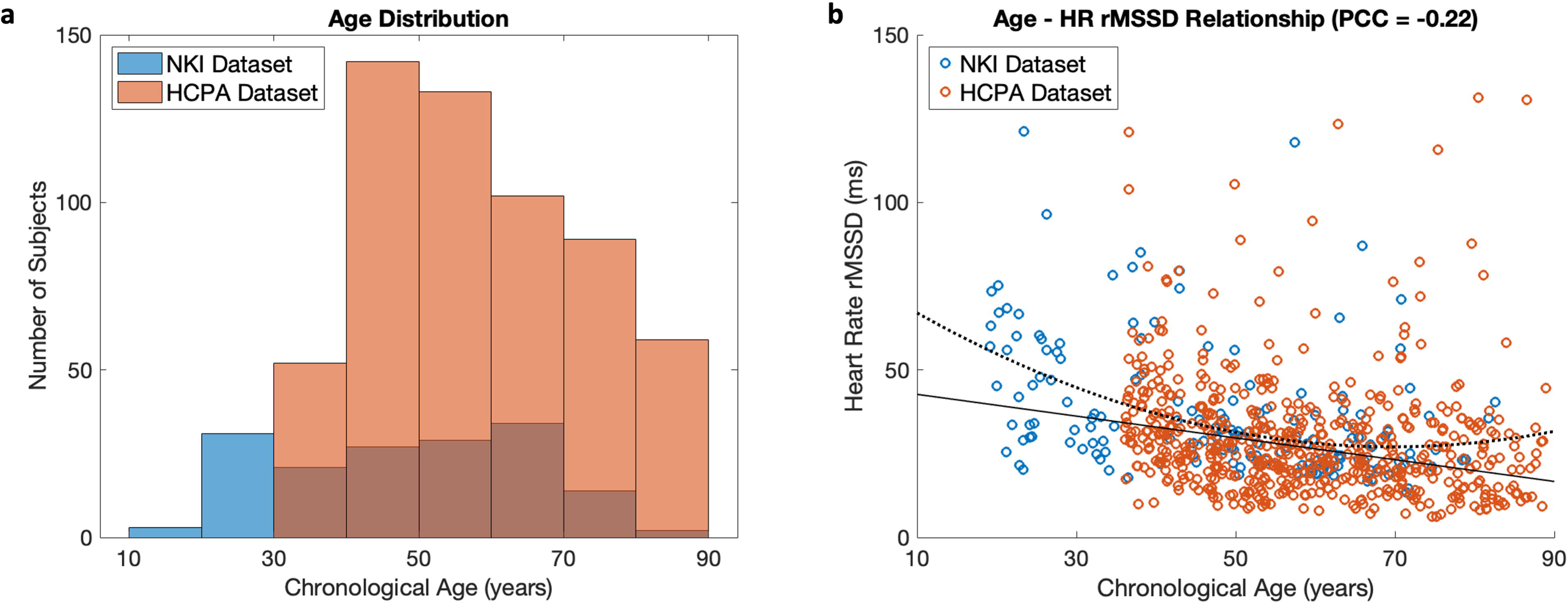
Age distribution of the NKI and HCPA datasets, and the relationship between age and heart rate variability (HRV) across the two datasets. a) Age distribution of the subjects included in the study. b) The relationship between age and HRV (measured as the root mean square of successive differences; rMSSD), together with fitted linear (solid line) and quadratic (dotted line) trends. For subjects with multiple segments, the HRV rMSSD measure was averaged across segments. PCC, Pearson correlation coefficient.

### Multivariate prediction of age from fMRI-physiological coupling

Distributed patterns of coupling between fMRI and low-frequency physiology were found to significantly predict age (**Table 1**, p < 0.05, permutation test). Age prediction significance was evaluated for NKI and HCPA datasets separately, and significant results mentioned in this section will refer to the case in which the performance on both datasets was significant based on permutation tests (p < 0.05). When using all 497 cortical, subcortical and white matter ROIs in the model, both linear and RBF kernels achieved significant age prediction results for the RV-, HR- and RVHR-fMRI variability. The second-order polynomial kernel, however, achieved significant age prediction results for both datasets only with the HR-fMRI variability.

**Table 1.**
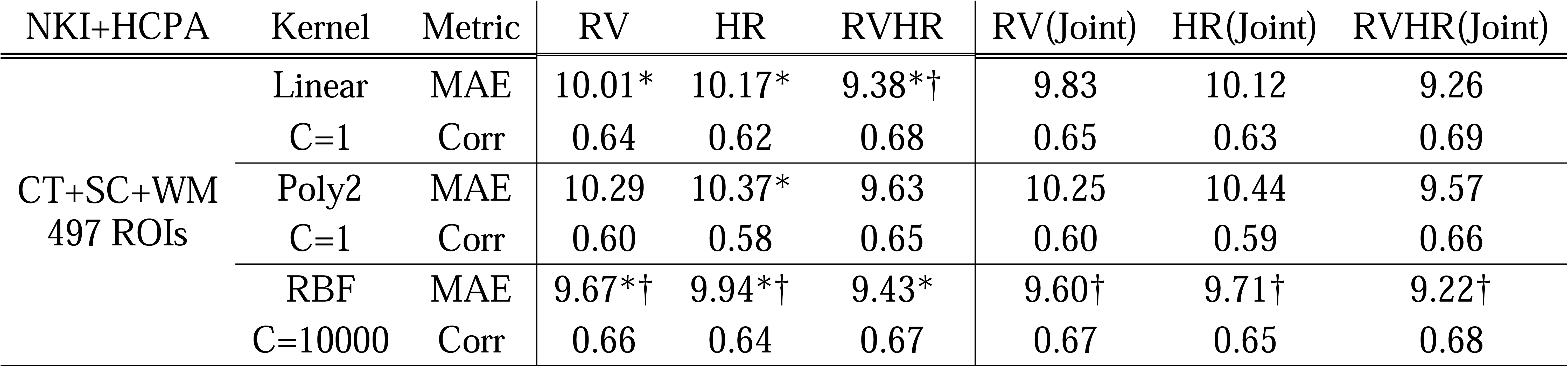
Age prediction performance with all 497 cortical (CT), subcortical (SC) and white matter (WM) ROIs across different SVR kernels (linear, 2nd degree polynomial (Poly2) and radial basis function (RBF)), fMRI physiological components (respiratory variation (RV), heart rate (HR), or RV and HR together (RVHR)), and training strategies. * indicates models that significantly outperformed permutation tests for both datasets when trained separately (p < 0.05; full p-values in **Table S2**). † marks the best age prediction performance (lowest mean absolute error; MAE, in years) for each column. Joint, training jointly on both NKI and HCPA datasets.

For all physiological components of the fMRI signal (RV-fMRI, HR-fMRI, and RVHR-fMRI variability), training jointly on NKI and HCPA datasets achieved better age prediction performance than when training on the two datasets separately. This was found to be the case regardless of which ROIs were used. Overall, the best performance was achieved using all 497 cortical, subcortical and white matter ROIs with a nonlinear RBF kernel (MAE_RVHR_ = 9.22 years), though the linear kernel achieved a very similar performance (MAE_RVHR_ = 9.26 years). The best prediction performance for RV- or HR-fMRI variability individually was achieved using all 497 ROIs and a nonlinear RBF kernel (**Fig. 4a-b**, MAE_RV_ = 9.60 years, MAE_HR_ = 9.71 years). The performance of different sets of ROIs in the jointly trained data, using RBF kernel, are shown in **Figure 5b**. Overall, adding subcortical and white matter ROIs to the set of cortical ROIs yielded gains in the accuracy of age prediction across training strategies and SVR kernels (**Figure 5b**; **Table S2**).

**Figure 4.**
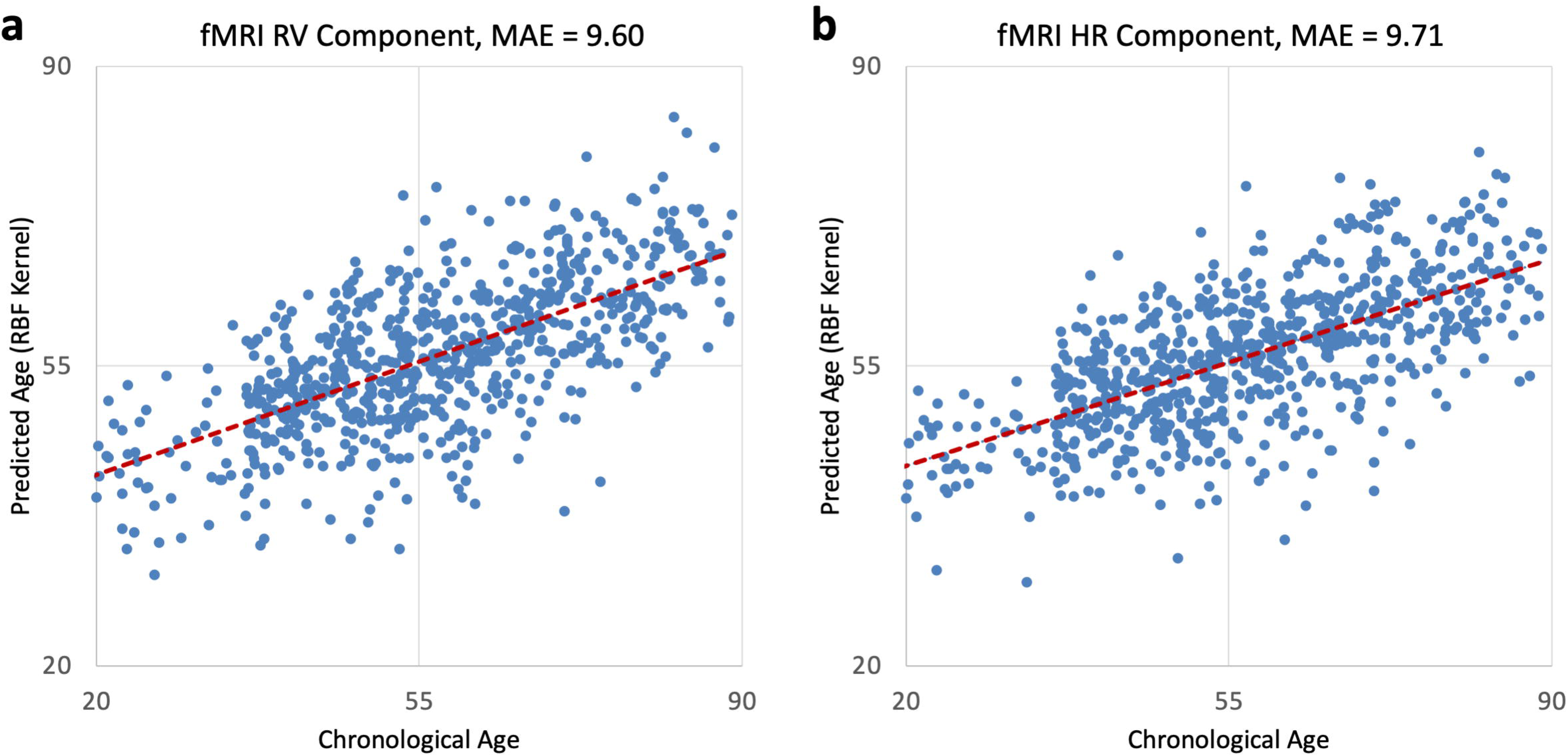
Scatter plot of the predicted age against the chronological age, shown here for the models attaining the best prediction performance using RV or HR-coupled fMRI variability. In both cases, the best performance was achieved using the RBF kernel and when training jointly across the two datasets (NKI, HCPA), as specified in **Table 1**. Age prediction accuracy is evaluated by mean absolute error (MAE, in years) between age and predicted age, averaged across all subjects with the 10 test folds concatenated.

**Figure 5.**
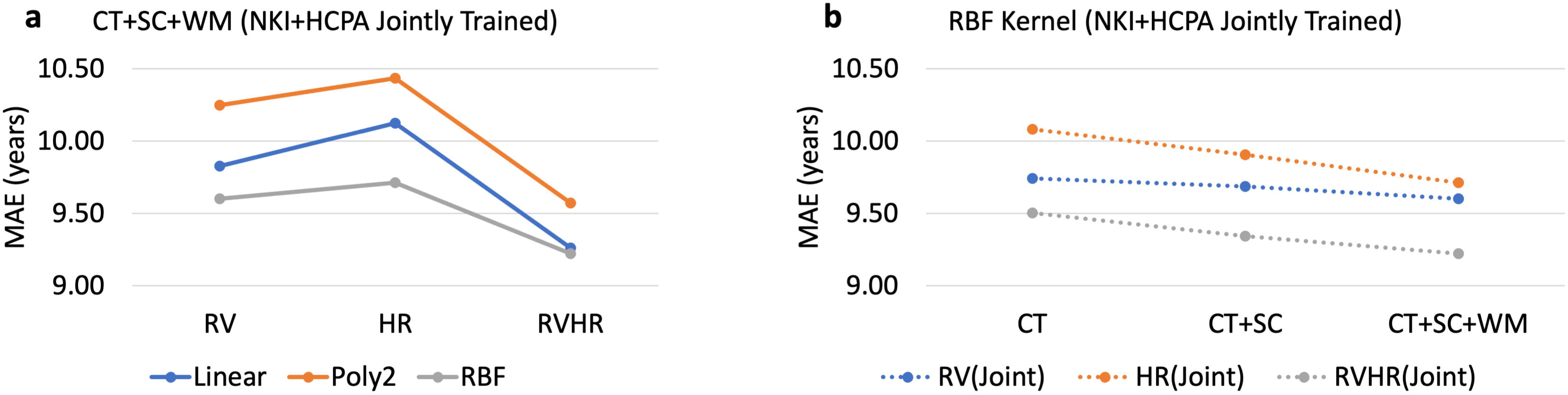
Effect of kernel selection and ROIs on the performance of age prediction from fMRI-physiological variability. Y-axis here is the mean absolute error (MAE, in years) between the chronological age and predicted age for the indicated (linear, 2nd degree polynomial (Poly2) and radial basis function (RBF)), fMRI physiological components (respiratory variation (RV), heart rate (HR), or RV and HR together (RVHR)) or ROI selections (cortical - CT, subcortical - SC and white matter - WM ROIs).

When comparing the age prediction performance across three kernels with different levels of nonlinearity, the nonlinear RBF kernel outperformed the linear and polynomial kernels consistently for models using either RV-fMRI or HR-fMRI variability independently (**Fig. 5a**; **Table S2**). The linear kernel was able to achieve similar or better age prediction results compared to the RBF kernel when the RVHR-fMRI variability was used as input. The Poly2 kernel performed the worst on almost all accounts, regardless of the input and training strategy. Detailed discussion on the impact of model nonlinearities and regularization are provided in **Text S1** (PedregosaFabian et al., 2011; Schulz et al., 2020; Tian et al., 2023; Bolt et al., 2025).

To test for a potential effect of age-related anatomic differences in the alignment to MNI space, we also constructed models in which the same 400 cortical ROIs were derived from either the MNI152-registered data or from the parcellation of individual subject anatomic images. Models using the subject-specific anatomic regions were found to achieve significant age prediction results compared to the permutation tests for the RV-fMRI variability with linear or RBF kernels, and for the RVHR-fMRI variability with the RBF kernel (**Table S3**). Nonetheless, ROIs derived from MNI152 template outperformed those derived from the individual T1-based ROIs across all SVR kernels; notably, this was also found to be the case when BOLD variability calculated from the raw and regressed signals, as described below. A detailed discussion of the potential role of age-related brain anatomy changes is provided in **Text S2**.

### Generalization to an independent dataset

We then tested the performance of age prediction from physio-fMRI variability on a new independent dataset to see if the age prediction model was generalizable to an unseen dataset. Models that were trained on the NKI dataset, or jointly on the NKI and HCPA datasets, were applied directly to the HRV-ER dataset; models trained on HCPA were not considered since this dataset does not encompass the age range of the younger adult group in the HRV-ER dataset. Here, only the HR-fMRI variability was tested since most of the respiratory belt recordings for the HRV-ER dataset were not of sufficient quality. The age prediction model trained on the NKI dataset yielded significantly higher predicted ages for the older versus younger adult groups (p < 0.005) in the HRV-ER dataset, where the best performance was achieved when the model was trained on the set of cortical and subcortical ROIs (**Table S4**). When jointly training on both NKI and HCPA datasets, cortical and subcortical ROIs combined also produced significantly different age prediction results between the two age groups (p < 0.005).

### Important brain networks and regions for age prediction with physiological-fMRI variability

To gain insight into the contribution of different large-scale functional brain networks to the prediction of age from physio-fMRI variability, we first tested the performance of models constructed from individual networks (**Fig. 6**; **Table S5**). For each model, only the ROIs corresponding to one of 17 cortical networks were used as input. When using RV-fMRI variability as input, one of the salience ventral attention networks (*SalVenAttnA*) achieved the best age prediction performance (MAE = 11.37 years). For the HR-fMRI variability, one of the somatomotor networks showed the best performance (*SomMotB;* MAE = 11.74 years). When fitting RV and HR jointly to the fMRI data, the networks that emerged when fitted separately showed the highest performance. One of the default mode networks showed one of the highest age prediction performances as well (*DefaultA;* MAE = 11.43 years).

**Figure 6.**
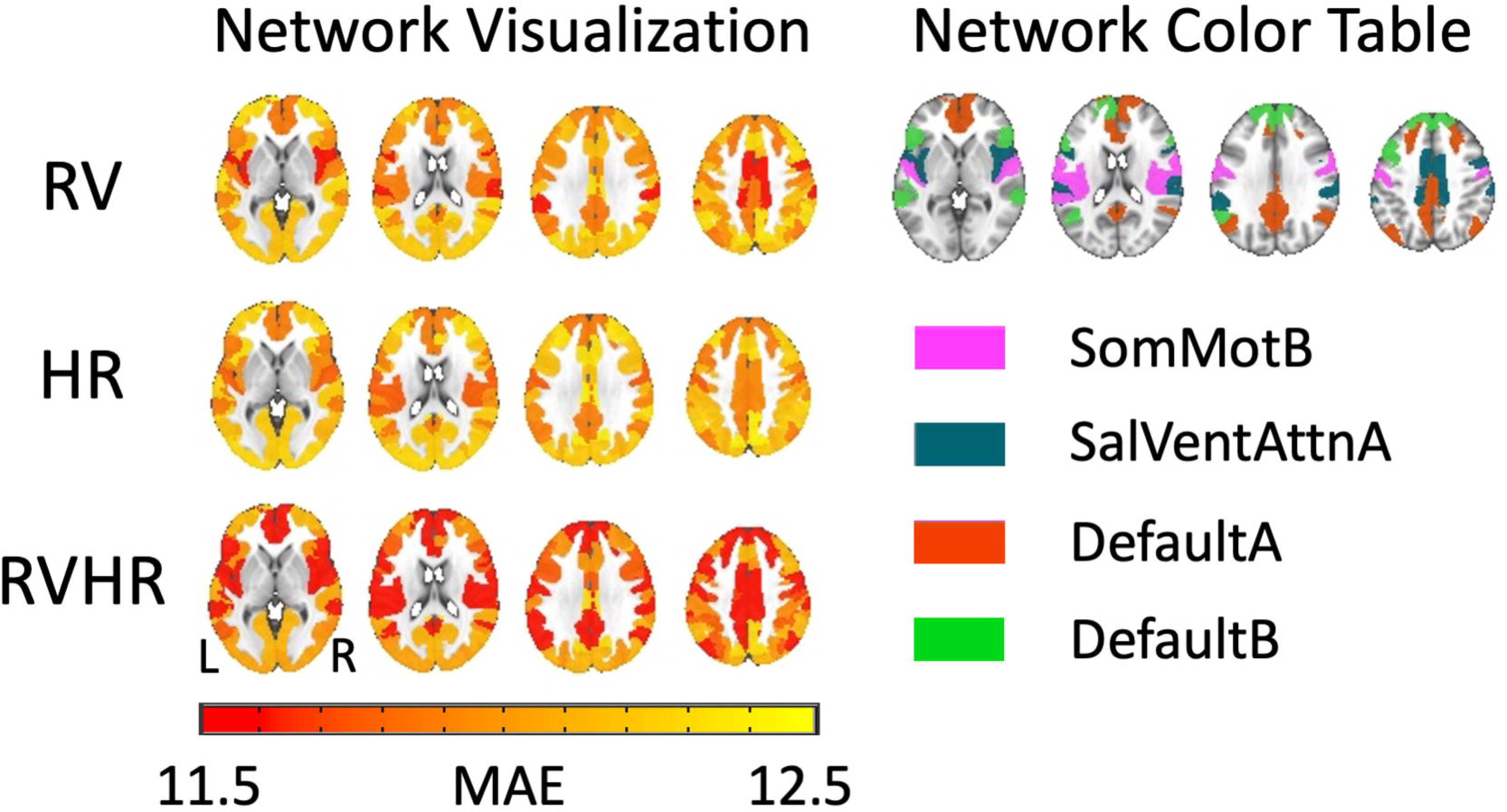
Network-level age prediction performance with radial basis function (RBF) kernel (C=1 based on the best validation set result). Parts of SalVenAttn, SomMot and Default networks were some of the strongest age predictors on the network level, as highlighted in red/orange exhibiting better age prediction accuracy or lower age prediction mean absolute error (MAE; in years). SalVenAttn - salience/ventral attention network; SomMot - somatomotor network; Default - default mode network. RV, respiratory variation coupled fMRI variability; HR, heart rate coupled fMRI variability; RVHR, RV and HR jointly coupled fMRI variability.

To further probe the importance of individual ROIs for age prediction, we also used SHapley Additive exPlanations (SHAP) analysis to examine how each ROI impacts age prediction output (Lundberg and Lee, 2017) (**Fig. 7**). When examining cortical ROIs, we found that bilateral insula, somatomotor cortex, and higher-order cortical regions were important for the prediction of age using the RV-fMRI variability, while the bilateral extrastriate cortex, somatomotor cortex and left lateral prefrontal cortex were important for age prediction using the HR-fMRI variability. When examining subcortical ROIs, the right hippocampus was the only region surviving this threshold consistently across both datasets for HR-fMRI variability, and was also present in the thresholded RV-fMRI maps. Left anterior thalamus and right posterior thalamus were the other regions passing this threshold for the RV-fMRI variability.

**Figure 7.**
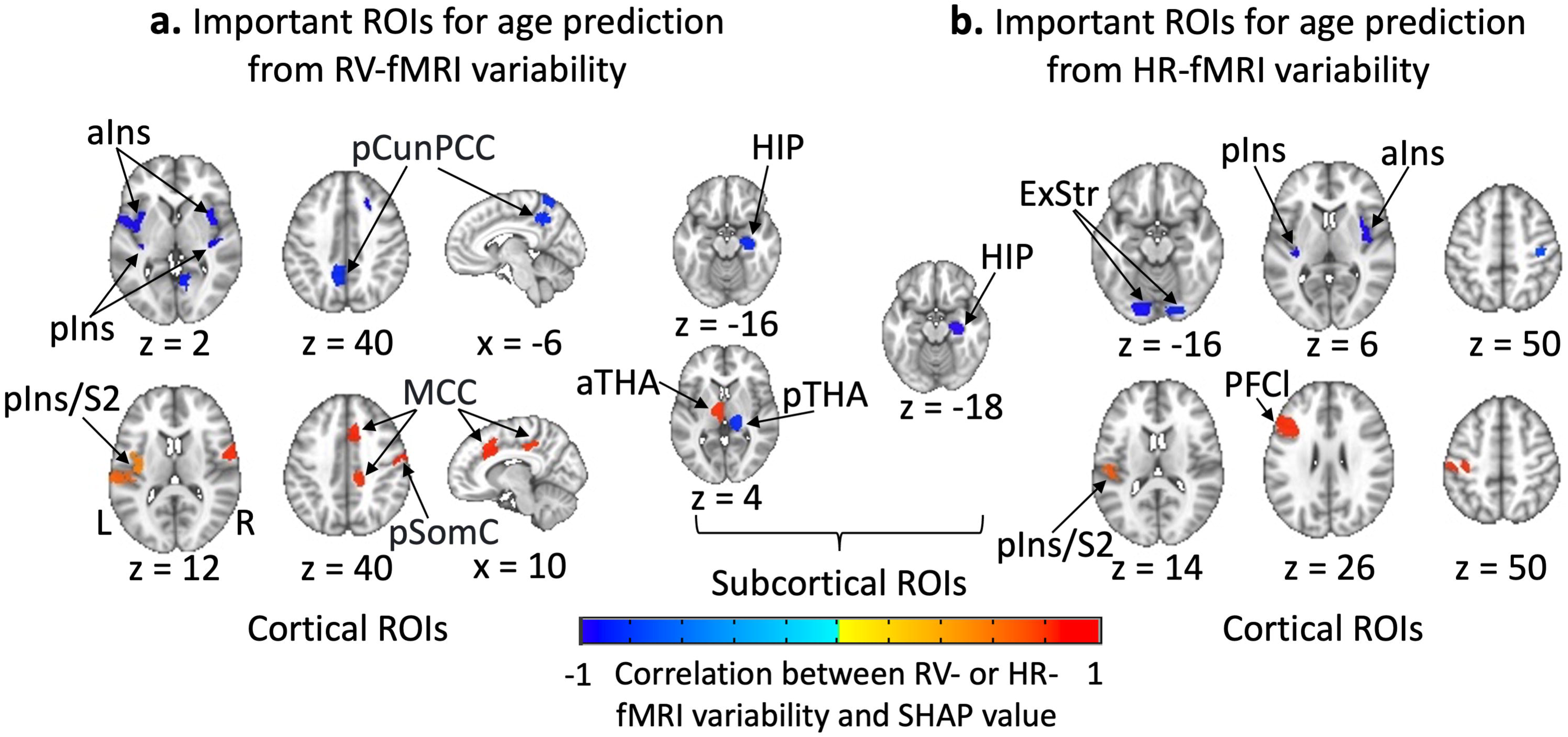
Important cortical and subcortical ROIs for predicting chronological age using the variability of a) RV-fitted or b) HR-fitted fMRI component, as evaluated by SHAP analysis. The models in this analysis used the RBF kernel and were trained separately on the NKI and HCPA datasets. Brain areas colored on the figure are those within the intersection of the top 20% most important ROIs across these two datasets. The color indicates the correlation value between the RV- or HR-fMRI variability for each of these ROIs and their corresponding SHAP values when trained jointly with both NKI and HCPA datasets. Positive values (red) indicate that a larger feature value (physiologically coupled fMRI variability) contributes to an older age prediction, and negative values (blue) indicate that a smaller fMRI-physiological variability value leads to an increase in age prediction. Primary somatosensory cortex - pSomC, secondary somatosensory cortex - S2, anterior insula - aIns, posterior insula - pIns, dorsal prefrontal cortex - PFCd, superior parietal lobule - SPL, post central - PostC, frontal operculum - FrOper, precuneus posterior cingulate cortex - pCunPCC, central cortex - Cent, extra-striate inferior - ExStrInf, parietal medial - ParMed, middle cingulate cortex - MCC, lateral prefrontal cortex - PFCl, extrastriate cortex - ExStr, extra-striate inferior - ExStrInf, hippocampus - HIP, anterior thalamus - aTHA, posterior thalamus - pTHA.

### Multivariate prediction of age from BOLD variability, before and after regressing out physiological components

For direct comparison with the wide literature on relationships between fMRI variability and aging, we also assessed the predictive of the non-z-normalized ‘raw’ fMRI variability (RAW), and the impact of regressing out low-frequency physiological (RV, HR) components (REG; **Table S6**). We observed that regressing out physiological signals did not reduce the prediction performance of BOLD variability measures, and in some cases led to improved performance. The best age prediction performance was achieved using ALFF after regressing out the joint RVHR component, when trained separately using RBF kernel and all 497 cortical, subcortical and white matter ROIs (MAE = 7.79 years).

Similar to the case of the fMRI-physiological variability, we observed that MNI-based parcellations outperformed T1-based cortical ROIs for both the RAW or REG input data (**Table S3**). Interestingly, unlike the case for the physiologically coupled fMRI variability, models trained on the two datasets combined did not always outperform the case of training separately.

## Discussion

This study aimed to determine whether brain-wide, multivariate patterns of low-frequency fMRI-physiology coupling can reliably predict chronological age in a community sample. Our goal was not to outperform alternative age prediction methods, but to test if the low-frequency physiological component of fMRI data is capable of predicting age, and to identify key contributing brain regions. We demonstrated that fMRI variability coupled to spontaneous respiratory volume and heart rate fluctuations was capable of predicting age beyond chance. Brain areas implicated in central autonomic regulation were amongst the strongest contributors to age prediction, including one of the salience/ventral attention networks comprising bilateral insula and part of the middle cingulate cortex. Furthermore, removing the fMRI physiological components improved the performance of age prediction from BOLD variability. These results indicate that 1) the coupling between fMRI and low-frequency physiological processes contains age-related information, and 2) relationships between aging and BOLD variability seen in the literature are not fully driven by changes in systemic physiology.

Overall, these results suggest that interactions between low-frequency autonomic processes and fMRI signals offer a unique non-invasive and high-resolution opportunity to study brain-body physiological coupling in the context of aging. These findings also open new avenues for designing biomarkers for the deterioration of cardiorespiratory or cerebrovascular health and their impact on brain health.

### Age prediction from low-frequency coupling between fMRI and autonomic physiology

We observed that the low-frequency physiological component of fMRI data can reliably predict age, and that its predictability does not solely arise from age-related differences in respiratory, cardiac, or fMRI signal characteristics themselves, but from the coupling between them. To test this, we performed permutation tests in which randomly selected, slow-varying physiological signals were fitted to the fMRI signals. Interestingly, we observed cases where the empirical results did not surpass the p<0.05 threshold compared with the permutation tests when predicting the age associated with (mismatched) fMRI data, suggesting that fMRI temporal and frequency profiles differ intrinsically with age. Nonetheless, when the empirical fMRI-physiological coupling was significant with respect to this null distribution, we may infer that age-related information in the coupling between fMRI and physiological signals exceeded that of either signal alone.

### Generalizability to an independent test dataset

The NKI, HCPA and HRV-ER datasets differ in MRI data acquisition parameters (e.g. TR, spatial resolution, and single- versus multi-echo sequences), physiological recording systems, preprocessing pipelines (e.g., FIX-ICA or ME-ICA), and age range, all of which might influence age prediction model generalizability. Despite these differences, a model trained on NKI and HCPA data obtained significantly different age predictions between younger and older adults in HRV-ER dataset, suggesting the robustness of physio-fMRI variability as an aging biomarker.

We did not include any harmonization techniques (Eshaghzadeh Torbati et al., 2021), and future work could assess how these approaches might improve the reliability and generalizability of this biomarker.

### Impact of removing physiological components on the relationship between age and BOLD variability

Removing the physiological component from the fMRI data did not remove the predictive ability of BOLD variability (**Table S6**), suggesting that intrinsic neural activity associated with processes other than autonomic or vascular responses also change robustly with age, and may reflect age-related reductions in dynamic range and cognitive flexibility (Garrett et al., 2013a; Uddin, 2021). Notably, fMRI ALFF after regressing out the RVHR component achieved the lowest mean absolute error for age prediction. Therefore, not only does the RVHR component itself convey information about aging, but disentangling RVHR from the remaining fMRI fluctuations (which putatively capture local neuronal activity) can help enhance the sensitivity of the latter to age-related neural changes. Thus, physio-fMRI coupling may serve a dual role: providing meaningful signal while probing brain-body interactions, but acting as confounds when the goal is to isolate neuronal activity.

Prior studies reported that subject-level cardiovascular and cerebrovascular indices together accounted for age effects in resting-state fluctuation amplitude (Tsvetanov et al., 2021a, 2021b), whereas our study focused on fMRI variations associated with low-frequency respiratory and cardiac fluctuations. This latter fMRI physiological component likely also contains neuronal regulation of autonomic processes, which is not captured in either the cerebrovascular or cardiovascular indices. Future studies could directly contrast these methods and compare the similarities or differences in what they convey about the aging process.

### Potential mechanisms behind age-related changes in fMRI-physiological coupling patterns

We would like to expand on two potential mechanisms underlying the observed age-related changes in fMRI-physiology coupling. One factor may stem from decreased cerebral vascular reactivity in older adults across both grey and white matter (Handwerker et al., 2007; Lu et al., 2011; Thomas et al., 2014; McKetton et al., 2018; Kapoor et al., 2025), which might limit the cerebrovascular response to low-frequency cardiac or respiratory variations. Relatedly, smaller BOLD responses(seen with both fMRI (Handwerker et al., 2007; Friedman et al., 2008) and functional near-infrared spectroscopy (fNIRS) data (Karunakaran et al., 2021)), reduced fMRI activation volume during breath-holding (Kannurpatti et al., 2010) have been reported. Hypercapnia studies conducted in both rodents (Balbi et al., 2015) and humans (Riecker et al., 2003; Flück et al., 2014) have found impaired vasodilation in older adults, consistent with age- related vessel stiffening (Yabluchanskiy et al., 2021) and nitric oxide endothelial pathway dysfunction, a mechanism thought to contribute to the coupling between fMRI BOLD signal and slow respiratory changes (Mayhan et al., 1990; Ungvari et al., 2008).

Second, aging has been associated with an imbalanced autonomic nervous system (ANS), with increased sympathetic nervous system (SNS) activity and decreased parasympathetic nervous system (PNS) activity (Abboud, 2010; Thayer et al., 2010; Giunta et al., 2024; Mather, 2024). Accordingly, older age in both NKI and HCPA was correlated with lower heart rate rMSSD (**Fig. 3b**), a marker of vagally mediated PNS activity that tends to decline with age (Thomas et al., 2019; Gullett et al., 2023). Interestingly, brain regions contributing most strongly to age prediction overlapped with known autonomic regulation hubs (Beissner et al., 2013), including left posterior insula (pIns), right anterior insula (aIns), and middle cingulate cortex (MCC) (Beissner et al., 2013; **Fig. 7**). SNS-related regions (e.g., left pIns, MCC, right primary somatosensory cortex, and left medial anterior thalamus) showed positive correlation between feature values and SHAP values (i.e., greater variability is associated with increasing age), consistent with increased SNS activity in older adults. On the other hand, PNS-related regions (e.g., right aIns, bilateral dorsal anterior insula, right hippocampus, and precuneus / posterior cingulate cortex) were negatively correlated with SHAP values, consistent with decreased PNS activity in older adults. Incorporating white matter regions into the model also improved age prediction accuracy, and since white matter fMRI signals are closely tied to SNS activity (Özbay et al., 2019; Picchioni et al., 2022), these findings may reflect age-related SNS alterations. Aside from ANS imbalance, the right anterior insular cortex has also been implicated in integrating autonomic signals with conscious thought processes, which might be compromised in older adults (Craig, 2002; Critchley et al., 2002; Critchley, 2004; Uddin, 2015).

### Limitations and future directions

One limitation of the current study is that other physiological or phenotypic covariates known to influence brain-body relationships with age (e.g. vigilance state, blood pressure; (Tucsek et al., 2014; Kimmerly, 2017)) are not modeled. Future studies could incorporate these features when modeling the brain-body relationship. Another limitation was the use of a physiological response function basis set developed on healthy young adults. Although the basis set allows flexibility when fitting RV or HR signals across ROIs or subjects, it might not accurately capture physio-fMRI coupling in older adults or those with pathological cardiovascular or cerebrovascular changes (O’Rourke and Hashimoto, 2007). Furthermore, we focused on the standard deviation of the fitted physiological components, while other variability metrics (e.g. rMSSD) and normalization schemes might reveal complementary aspects of physio-fMRI coupling. In addition, the 62.5 Hz PPG sampling rate in the NKI dataset might not be as reliable for detecting heart rate; the minimal sampling frequency for PPG recording to accurately detect HRV varies in the literature, ranging from 25 Hz to 250 Hz (Kwon et al., 2018; Béres and Hejjel, 2021). Despite our efforts to control for head motion, residual head motion may still affect fMRI physiological components and age prediction performance. Initial explorations incorporating head motion as an additional feature in age prediction models (**Text S3, Table S7)** showed marginal performance improvements.

Beyond methodological considerations, future studies could examine how physio-fMRI-based brain age relates to cognitive or behavioral performance; our preliminary exploration and results are presented in **Text S4 & Table S8** (Cumplido-Mayoral et al., 2023; Grady et al., 2023; Gaser et al., 2024; Goodman et al., 2024; Kim et al., 2025). Further, the understanding of aging-related fMRI-physiology coupling changes might be translatable to diseases that share similar mechanisms as normal aging, or have risk profiles that are related to accelerated cerebral vascular aging (e.g., diabetes, obesity, or hypertension). The current prediction framework for modeling brain-body interactions could be extended from healthy aging to patient populations with cognitive impairment, dementia, or cardiorespiratory health problems.

## Acknowledgments

This work was supported by NIH RF1MH125931 (C.C., M.M.), T32MH064913 (K.R.), F99AG079810 (S.E.G.), and T32EB021937 (H.P.), and the Sally and Dave Hopkins Faculty Fellowship (C.C.). LQU is supported by the National Institute of Child Health and Human Development (R21HD111805 and R01HD11669) and the National Institute on Drug Abuse (U24DA041147 and U01DA050987).

## References

Abboud FM (2010) The Walter B. Cannon Memorial Award Lecture, 2009. Physiology in perspective: The wisdom of the body. In search of autonomic balance: the good, the bad, and the ugly. Am J Physiol Regul Integr Comp Physiol 298:R1449–R1467.

Avants BB, Tustison NJ, Song G, Cook PA, Klein A, Gee JC (2011) A reproducible evaluation of ANTs similarity metric performance in brain image registration. Neuroimage 54:2033–2044.

Balbi M, Ghosh M, Longden TA, Jativa Vega M, Gesierich B, Hellal F, Lourbopoulos A, Nelson MT, Plesnila N (2015) Dysfunction of mouse cerebral arteries during early aging. J Cereb Blood Flow Metab 35:1445–1453.

Bayrak RG, Hansen CB, Salas JA, Ahmed N, Lyu I, Mather M, Huo Y, Chang C (2025) DeepPhysioRecon: Tracing peripheral physiology in low frequency fMRI dynamics. Imaging Neurosci (Camb) 3:IMAG. a. 163.

Beissner F, Meissner K, Bär K-J, Napadow V (2013) The autonomic brain: an activation likelihood estimation meta-analysis for central processing of autonomic function. J Neurosci 33:10503–10511.

Béres S, Hejjel L (2021) The minimal sampling frequency of the photoplethysmogram for accurate pulse rate variability parameters in healthy volunteers. Biomed Signal Process Control 68:102589.

Birn RM, Diamond JB, Smith MA, Bandettini PA (2006) Separating respiratory-variation-related fluctuations from neuronal-activity-related fluctuations in fMRI. Neuroimage 31:1536–1548.

Biswal B, Yetkin FZ, Haughton VM, Hyde JS (1995) Functional connectivity in the motor cortex of resting human brain using echo-planar MRI. Magn Reson Med 34:537–541.

Bolt T, Wang S, Nomi JS, Setton R, Gold BP, deB Frederick B, Yeo BTT, Chen JJ, Picchioni D, Duyn JH, Spreng RN, Keilholz SD, Uddin LQ, Chang C (2025) Autonomic physiological coupling of the global fMRI signal. Nat Neurosci:1–9.

Bookheimer SY et al. (2019) The Lifespan Human Connectome Project in Aging: An overview. Neuroimage 185:335–348.

Boylan MA, Foster CM, Pongpipat EE, Webb CE, Rodrigue KM, Kennedy KM (2021) Greater BOLD Variability is Associated With Poorer Cognitive Function in an Adult Lifespan Sample. Cereb Cortex 31:562–574.

Bright MG, Whittaker JR, Driver ID, Murphy K (2020) Vascular physiology drives functional brain networks. Neuroimage 217:116907.

Chang C, Cunningham JP, Glover GH (2009) Influence of heart rate on the BOLD signal: The cardiac response function. Neuroimage 44:857–869.

Chen JE, Lewis LD, Chang C, Tian Q, Fultz NE, Ohringer NA, Rosen BR, Polimeni JR (2020) Resting-state “physiological networks.” Neuroimage 213:116707.

Chen JJ (2019) Functional MRI of brain physiology in aging and neurodegenerative diseases. Neuroimage 187:209–225.

Craig AD (2002) How do you feel? Interoception: the sense of the physiological condition of the body. Nat Rev Neurosci 3:655–666.

Critchley HD (2004) The human cortex responds to an interoceptive challenge. Proc Natl Acad Sci U S A 101:6333–6334.

Critchley HD, Melmed RN, Featherstone E, Mathias CJ, Dolan RJ (2002) Volitional control of autonomic arousal: a functional magnetic resonance study. Neuroimage 16:909–919.

Cumplido-Mayoral I et al. (2023) Biological brain age prediction using machine learning on structural neuroimaging data: Multi-cohort validation against biomarkers of Alzheimer’s disease and neurodegeneration stratified by sex. Elife 12:e81067.

Dale AM, Fischl B, Sereno MI (1999) Cortical Surface-Based Analysis. Neuroimage 9:179–194.

Delli Pizzi S, Gambi F, Di Pietro M, Caulo M, Sensi SL, Ferretti A (2023) BOLD cardiorespiratory pulsatility in the brain: from noise to signal of interest. Front Hum Neurosci 17:1327276.

Duyn JH, Ozbay PS, Chang C, Picchioni D (2020) Physiological changes in sleep that affect fMRI inference. Curr Opin Behav Sci 33:42–50.

Edlow BL, Takahashi E, Wu O, Benner T, Dai G, Bu L, Grant PE, Greer DM, Greenberg SM, Kinney HC, Folkerth RD (2012) Neuroanatomic Connectivity of the Human Ascending Arousal System Critical to Consciousness and Its Disorders. J Neuropathol Exp Neurol 71:531–546.

Egorova N, Veldsman M, Cumming T, Brodtmann A (2017) Fractional amplitude of low-frequency fluctuations (fALFF) in post-stroke depression. NeuroImage Clin 16:116–124.

Eshaghzadeh Torbati M, Minhas DS, Ahmad G, O’Connor EE, Muschelli J, Laymon CM, Yang Z, Cohen AD, Aizenstein HJ, Klunk WE, Christian BT, Hwang SJ, Crainiceanu CM, Tudorascu DL (2021) A multi-scanner neuroimaging data harmonization using RAVEL and ComBat. Neuroimage 245:118703.

Fan J, Juttukonda MR, Goodale SE, Wang S, Orbán C, Varadarajan D, Polimeni JR, Chang C, Salat DH, Chen JE (2025) Functional MRI signatures of autonomic physiology in aging. Commun Biol 8:1287.

Fischl B, Salat DH, Busa E, Albert M, Dieterich M, Haselgrove C, van der Kouwe A, Killiany R, Kennedy D, Klaveness S, Montillo A, Makris N, Rosen B, Dale AM (2002) Whole Brain Segmentation. Neuron 33:341–355.

Fischl B, Sereno MI, Dale AM (1999) Cortical Surface-Based Analysis. Neuroimage 9:195–207.

Flück D, Beaudin AE, Steinback CD, Kumarpillai G, Shobha N, McCreary CR, Peca S, Smith EE, Poulin MJ (2014) Effects of aging on the association between cerebrovascular responses to visual stimulation, hypercapnia and arterial stiffness. Front Physiol 5:49.

Friedman L, Turner JA, Stern H, Mathalon DH, Trondsen LC, Potkin SG (2008) Chronic smoking and the BOLD response to a visual activation task and a breath hold task in patients with schizophrenia and healthy controls. Neuroimage 40:1181–1194.

Garrett DD, Kovacevic N, McIntosh AR, Grady CL (2010) Blood Oxygen Level-Dependent Signal Variability Is More than Just Noise. J Neurosci 30:4914–4921.

Garrett DD, Kovacevic N, McIntosh AR, Grady CL (2013a) The Modulation of BOLD Variability between Cognitive States Varies by Age and Processing Speed. Cereb Cortex 23:684–693.

Garrett DD, Samanez-Larkin GR, MacDonald SWS, Lindenberger U, McIntosh AR, Grady CL (2013b) Moment-to-moment brain signal variability: A next frontier in human brain mapping? Neurosci Biobehav Rev 37:610–624.

Gaser C, Kalc P, Cole JH (2024) A perspective on brain-age estimation and its clinical promise. Nat Comput Sci 4:744–751.

Giunta S, Xia S, Pelliccioni G, Olivieri F (2024) Autonomic nervous system imbalance during aging contributes to impair endogenous anti-inflammaging strategies. GeroScience 46:113–127.

Glasser MF, Sotiropoulos SN, Wilson JA, Coalson TS, Fischl B, Andersson JL, Xu J, Jbabdi S, Webster M, Polimeni JR, Van Essen DC, Jenkinson M (2013) The minimal preprocessing pipelines for the Human Connectome Project. Neuroimage 80:105–124.

Glover GH, Li T-Q, Ress D (2000) Image-based method for retrospective correction of physiological motion effects in fMRI: RETROICOR. Magn Reson Med 44:162–167.

Goodman ZT, Nomi JS, Kornfeld S, Bolt T, Saumure RA, Romero C, Bainter SA, Uddin LQ (2024) Brain signal variability and executive functions across the life span. Netw Neurosci 8:226–240.

Grady CL, Garrett DD (2014) Understanding variability in the BOLD signal and why it matters for aging. Brain Imaging Behav 8:274–283.

Grady CL, Rieck JR, Baracchini G, DeSouza B (2023) Relation of resting brain signal variability to cognitive and socioemotional measures in an adult lifespan sample. Soc Cogn Affect Neurosci 18 Available at: 10.1093/scan/nsad044.

Guitart-Masip M, Salami A, Garrett D, Rieckmann A, Lindenberger U, Bäckman L (2016) BOLD Variability is Related to Dopaminergic Neurotransmission and Cognitive Aging. Cereb Cortex 26:2074–2083.

Gullett N, Zajkowska Z, Walsh A, Harper R, Mondelli V (2023) Heart rate variability (HRV) as a way to understand associations between the autonomic nervous system (ANS) and affective states: A critical review of the literature. Int J Psychophysiol 192:35–42.

Handwerker DA, Gazzaley A, Inglis BA, D’Esposito M (2007) Reducing vascular variability of fMRI data across aging populations using a breathholding task. Hum Brain Mapp 28:846–859.

Hansen CB, Yang Q, Lyu I, Rheault F, Kerley C, Chandio BQ, Fadnavis S, Williams O, Shafer AT, Resnick SM, Zald DH, Cutting LE, Taylor WD, Boyd B, Garyfallidis E, Anderson AW, Descoteaux M, Landman BA, Schilling KG (2021) Pandora: 4-D White Matter Bundle Population-Based Atlases Derived from Diffusion MRI Fiber Tractography. Neuroinformatics 19:447–460.

Harms MP et al. (2018) Extending the Human Connectome Project across ages: Imaging protocols for the Lifespan Development and Aging projects. Neuroimage 183:972–984.

Hu S, Chao HH-A, Zhang S, Ide JS, Li C-SR (2014) Changes in cerebral morphometry and amplitude of low-frequency fluctuations of BOLD signals during healthy aging: correlation with inhibitory control. Brain Struct Funct 219:983–994.

Kannurpatti SS, Motes MA, Rypma B, Biswal BB (2010) Neural and vascular variability and the fMRI-BOLD response in normal aging. Magn Reson Imaging 28:466–476.

Kapoor A, Dutt S, Engstrom AC, Alitin JPM, Lohman T, Sible IJ, Marshall A, Shenasa F, Gaubert A, Bradford DR, Sordo L, Shao X, Rodgers K, Head E, Wang DJ, Nation DA (2025) Association of medial temporal lobe cerebrovascular reactivity and memory function in older adults with and without cognitive impairment. Neurology 104:e210210.

Karavallil Achuthan S, Coburn KL, Beckerson ME, Kana RK (2023) Amplitude of low frequency fluctuations during resting state fMRI in autistic children. Autism Res 16:84–98.

Karunakaran KD, Ji K, Chen DY, Chiaravalloti ND, Niu H, Alvarez TL, Biswal BB (2021) Relationship between age and cerebral hemodynamic response to breath holding: A functional near-infrared spectroscopy study. Brain Topogr 34:154–166.

Kassinopoulos M, Mitsis GD (2019) Identification of physiological response functions to correct for fluctuations in resting-state fMRI related to heart rate and respiration. Neuroimage 202:116150.

Kim H, Park S, Seo SW, Na DL, Jang H, Kim JP, Kim HJ, Kang SH, Kwak K (2025) A novel deep learning-based brain age prediction framework for routine clinical MRI scans. NPJ Aging 11:70.

Kimmerly DS (2017) A review of human neuroimaging investigations involved with central autonomic regulation of baroreflex-mediated cardiovascular control. Auton Neurosci 207:10–21.

Kumral D, Schaare HL, Beyer F, Reinelt J, Uhlig M, Liem F, Lampe L, Babayan A, Reiter A, Erbey M, Roebbig J, Loeffler M, Schroeter ML, Husser D, Witte AV, Villringer A, Gaebler M (2019) The age-dependent relationship between resting heart rate variability and functional brain connectivity. Neuroimage 185:521–533.

Kundu P, Inati SJ, Evans JW, Luh W-M, Bandettini PA (2012) Differentiating BOLD and non-BOLD signals in fMRI time series using multi-echo EPI. Neuroimage 60:1759–1770.

Kundu P, Voon V, Balchandani P, Lombardo MV, Poser BA, Bandettini PA (2017) Multi-echo fMRI: A review of applications in fMRI denoising and analysis of BOLD signals. Neuroimage 154:59–80.

Kwon O, Jeong J, Kim HB, Kwon IH, Park SY, Kim JE, Choi Y (2018) Electrocardiogram sampling frequency range acceptable for heart rate variability analysis. Healthc Inform Res 24:198–206.

Li M, Gao Y, Lawless RD, Xu L, Zhao Y, Schilling KG, Ding Z, Anderson AW, Landman BA, Gore JC (2023) Changes in white matter functional networks across late adulthood. Front Aging Neurosci 15:1204301.

Li M, Schilling KG, Gao F, Xu L, Choi S, Gao Y, Zu Z, Anderson AW, Ding Z, Landman BA, Gore JC (2024) Quantification of mediation effects of white matter functional characteristics on cognitive decline in aging. Cereb Cortex 34:bhae114.

Lu H, Xu F, Rodrigue KM, Kennedy KM, Cheng Y, Flicker B, Hebrank AC, Uh J, Park DC (2011) Alterations in Cerebral Metabolic Rate and Blood Supply across the Adult Lifespan. Cereb Cortex 21:1426–1434.

Lundberg SM, Lee S-I (2017) A unified approach to interpreting model predictions. Neural Inf Process Syst:4765–4774.

Mather M (2024) The emotion paradox in the aging body and brain. Ann N Y Acad Sci 1536:13–41.

Mayhan WG, Faraci FM, Baumbach GL, Heistad DD (1990) Effects of aging on responses of cerebral arterioles. Am J Physiol 258:H1138–H1143.

McKetton L, Sobczyk O, Duffin J, Poublanc J, Sam K, Crawley AP, Venkatraghavan L, Fisher JA, Mikulis DJ (2018) The aging brain and cerebrovascular reactivity. Neuroimage 181:132–141.

Millar PR, Petersen SE, Ances BM, Gordon BA, Benzinger TLS, Morris JC, Balota DA (2020) Evaluating the Sensitivity of Resting-State BOLD Variability to Age and Cognition after Controlling for Motion and Cardiovascular Influences: A Network-Based Approach. Cereb Cortex 30:5686–5701.

Montalà-Flaquer M, Cañete-Massé C, Vaqué-Alcázar L, Bartrés-Faz D, Peró-Cebollero M, Guàrdia-Olmos J (2022) Spontaneous brain activity in healthy aging: An overview through fluctuations and regional homogeneity. Front Aging Neurosci 14:1002811.

Murphy K, Birn RM, Bandettini PA (2013) Resting-state fMRI confounds and cleanup. Neuroimage 80:349–359.

Nomi JS, Bolt TS, Ezie CEC, Uddin LQ, Heller AS (2017) Moment-to-Moment BOLD Signal Variability Reflects Regional Changes in Neural Flexibility across the Lifespan. J Neurosci 37:5539–5548.

Nomi JS, Bzdok D, Li J, Bolt T, Chang C, Kornfeld S, Goodman ZT, Yeo BTT, Spreng RN, Uddin LQ (2024) Systematic cross-sectional age-associations in global fMRI signal topography. Imaging Neuroscience 2:1–13.

Nooner KB et al. (2012) The NKI-Rockland Sample: A Model for Accelerating the Pace of Discovery Science in Psychiatry. Front Neurosci 6 Available at: 10.3389/fnins.2012.00152.

O’Rourke MF, Hashimoto J (2007) Mechanical factors in arterial aging: a clinical perspective. J Am Coll Cardiol 50:1–13.

Özbay PS, Chang C, Picchioni D, Mandelkow H, Chappel-Farley MG, van Gelderen P, de Zwart JA, Duyn J (2019) Sympathetic activity contributes to the fMRI signal. Commun Biol 2:421.

Özbay PS, Chang C, Picchioni D, Mandelkow H, Moehlman TM, Chappel-Farley MG, van Gelderen P, de Zwart JA, Duyn JH (2018) Contribution of systemic vascular effects to fMRI activity in white matter. Neuroimage 176:541–549.

PedregosaFabian, VaroquauxGaël, GramfortAlexandre, MichelVincent, ThirionBertrand, GriselOlivier, BlondelMathieu, PrettenhoferPeter, WeissRon, DubourgVincent, VanderplasJake, PassosAlexandre, CournapeauDavid, BrucherMatthieu, PerrotMatthieu, DuchesnayÉdouard (2011) Scikit-learn: Machine Learning in Python. J Mach Learn Res Available at: 10.5555/1953048.2078195 [Accessed December 1, 2025].

Picchioni D, Özbay PS, Mandelkow H, de Zwart JA, Wang Y, van Gelderen P, Duyn JH (2022) Autonomic arousals contribute to brain fluid pulsations during sleep. Neuroimage 249:118888.

Riecker A, Grodd W, Klose U, Schulz JB, Gröschel K, Erb M, Ackermann H, Kastrup A (2003) Relation between regional functional MRI activation and vascular reactivity to carbon dioxide during normal aging. J Cereb Blood Flow Metab 23:565–573.

Salimi-Khorshidi G, Douaud G, Beckmann CF, Glasser MF, Griffanti L, Smith SM (2014) Automatic denoising of functional MRI data: Combining independent component analysis and hierarchical fusion of classifiers. Neuroimage 90:449–468.

Schaefer A, Kong R, Gordon EM, Laumann TO, Zuo X-N, Holmes AJ, Eickhoff SB, Yeo BTT (2018) Local-Global Parcellation of the Human Cerebral Cortex from Intrinsic Functional Connectivity MRI. Cereb Cortex 28:3095–3114.

Schulz M-A, Yeo BTT, Vogelstein JT, Mourao-Miranada J, Kather JN, Kording K, Richards B, Bzdok D (2020) Different scaling of linear models and deep learning in UKBiobank brain images versus machine-learning datasets. Nat Commun 11:4238.

Song R, Min J, Wang S, Goodale SE, Rogge-Obando K, Yang R, Yoo HJ, Nashiro K, Chen JE, Mather M, Chang C (2025) The physiological component of the BOLD signal: Impact of age and heart rate variability biofeedback training. Imaging Neurosci (Camb) 3:IMAG.a.99.

Thayer JF, Yamamoto SS, Brosschot JF (2010) The relationship of autonomic imbalance, heart rate variability and cardiovascular disease risk factors. Int J Cardiol 141:122–131.

Thomas BL, Claassen N, Becker P, Viljoen M (2019) Validity of commonly used heart rate variability markers of autonomic nervous system function. Neuropsychobiology 78:14–26.

Thomas BP, Liu P, Park DC, van Osch MJP, Lu H (2014) Cerebrovascular reactivity in the brain white matter: magnitude, temporal characteristics, and age effects. J Cereb Blood Flow Metab 34:242–247.

Tian YE, Cropley V, Maier AB, Lautenschlager NT, Breakspear M, Zalesky A (2023) Heterogeneous aging across multiple organ systems and prediction of chronic disease and mortality. Nat Med 29:1221–1231.

Tian Y, Margulies DS, Breakspear M, Zalesky A (2020) Topographic organization of the human subcortex unveiled with functional connectivity gradients. Nat Neurosci 23:1421–1432.

Tsvetanov KA, Henson RNA, Jones PS, Mutsaerts H, Fuhrmann D, Tyler LK, Cam-CAN, Rowe JB (2021a) The effects of age on resting-state BOLD signal variability is explained by cardiovascular and cerebrovascular factors. Psychophysiology 58:e13714.

Tsvetanov KA, Henson RNA, Rowe JB (2021b) Separating vascular and neuronal effects of age on fMRI BOLD signals: Neurovascular ageing. Philosophical Transactions of the Royal Society B: Biological Sciences 376 Available at: 10.1098/rstb.2019.0631.

Tucsek Z, Toth P, Tarantini S, Sosnowska D, Gautam T, Warrington JP, Giles CB, Wren JD, Koller A, Ballabh P, Sonntag WE, Ungvari Z, Csiszar A (2014) Aging exacerbates obesity-induced cerebromicrovascular rarefaction, neurovascular uncoupling, and cognitive decline in mice. J Gerontol A Biol Sci Med Sci 69:1339–1352.

Tu W, Zhang N (2022) Neural underpinning of a respiration-associated resting-state fMRI network. Elife 11 Available at: 10.7554/eLife.81555.

Uddin LQ (2015) Salience processing and insular cortical function and dysfunction. Nat Rev Neurosci 16:55–61.

Uddin LQ (2020) Bring the noise: Reconceptualizing spontaneous neural activity. Trends Cogn Sci 24:734–746.

Uddin LQ (2021) Cognitive and behavioural flexibility: neural mechanisms and clinical considerations. Nat Rev Neurosci 22:167–179.

Ungvari Z, Labinskyy N, Gupte S, Chander PN, Edwards JG, Csiszar A (2008) Dysregulation of mitochondrial biogenesis in vascular endothelial and smooth muscle cells of aged rats. Am J Physiol Heart Circ Physiol 294:H2121–H2128.

Wang S, Rao B, Chen L, Chen Z, Fang P, Miao G, Xu H, Liao W (2021) Using fractional amplitude of low-frequency fluctuations and functional connectivity in patients with post-stroke cognitive impairment for a simulated stimulation program. Front Aging Neurosci 13:724267.

Waschke L, Kloosterman NA, Obleser J, Garrett DD (2021) Behavior needs neural variability. Neuron 109:751–766.

Wise RG, Ide K, Poulin MJ, Tracey I (2004) Resting fluctuations in arterial carbon dioxide induce significant low frequency variations in BOLD signal. Neuroimage 21:1652–1664.

Xu L, Gao Y, Li M, Lawless R, Zhao Y, Schilling KG, Rogers BP, Anderson AW, Ding Z, Landman BA, Gore JC (2024) Functional correlation tensors in brain white matter and the effects of normal aging. Brain Imaging Behav 18:1197–1214.

Yabluchanskiy A, Nyul-Toth A, Csiszar A, Gulej R, Saunders D, Towner R, Turner M, Zhao Y, Abdelkari D, Rypma B, Tarantini S (2021) Age-related alterations in the cerebrovasculature affect neurovascular coupling and BOLD fMRI responses: Insights from animal models of aging. Psychophysiology 58:e13718.

Yang L, Wei A-H, Ouyang T-T, Cao Z-Z, Duan A-W, Zhang H-H (2021) Functional plasticity abnormalities over the lifespan of first-episode patients with major depressive disorder: a resting state fMRI study. Ann Transl Med 9:349.

Yan L, Zhuo Y, Wang B, Wang DJJ (2011) Loss of coherence of low frequency fluctuations of BOLD FMRI in visual cortex of healthy aged subjects. Open Neuroimag J 5:105–111.

Yoo HJ et al. (2023) Multimodal neuroimaging data from a 5-week heart rate variability biofeedback randomized clinical trial. Sci Data 10:503.

Zang Y-F, He Y, Zhu C-Z, Cao Q-J, Sui M-Q, Liang M, Tian L-X, Jiang T-Z, Wang Y-F (2007) Altered baseline brain activity in children with ADHD revealed by resting-state functional MRI. Brain Dev 29:83–91.

Zhang X, Xue C, Cao X, Yuan Q, Qi W, Xu W, Zhang S, Huang Q (2021) Altered patterns of amplitude of low-frequency fluctuations and fractional amplitude of low-frequency fluctuations between amnestic and vascular mild cognitive impairment: An ALE-based comparative meta-analysis. Front Aging Neurosci 13:711023.

Zhong XZ, Chen JJ (2022) Resting-state functional magnetic resonance imaging signal variations in aging: The role of neural activity. Hum Brain Mapp 43:2880–2897.

Zou Q-H, Zhu C-Z, Yang Y, Zuo X-N, Long X-Y, Cao Q-J, Wang Y-F, Zang Y-F (2008) An improved approach to detection of amplitude of low-frequency fluctuation (ALFF) for resting-state fMRI: fractional ALFF. J Neurosci Methods 172:137–141.

