## Supplemental Table 1 for "Distributed fMRI patterns coupled to low-frequency cardiorespiratory dynamics provide markers of aging"

|  | Metric | RV | HR | RVHR |
| --- | --- | --- | --- | --- |
| NKI | Corr | 0.19 | 0.22 | 0.15 |
|  | p-value | 0.02 | <0.01 | 0.05 |
| HCPA | Corr | -0.01 | 0.01 | -0.07 |
|  | p-value | 0.85 | 0.87 | 0.11 |

**Table S1.** Correlation between mean framewise displacement and averaged physio-fMRI variability (respiratory variation (RV), heart rate (HR), or RV and HR together (RVHR)) across 497 ROIs, for the NKI and HCPA datasets. RV-fMRI variability and HR-fMRI variability in the NKI dataset are significantly correlated with head motion (p<0.05), but the relationships are not significant in the NKI dataset for RVHR-fMRI variability, or in the HCPA dataset for any physiological measure. Corr, Pearson correlation coefficient.
