## Supplemental Table 2 for "Distributed fMRI patterns coupled to low-frequency cardiorespiratory dynamics provide markers of aging"

| NKI+HCPA-MNI | Kernel | Metric | RV | | HR | | RVHR | | RV(Joint) | HR(Joint) | RVHR(Joint) |
| --- | --- | --- | --- | --- | --- | --- | --- | --- | --- | --- | --- |
| CT  400 ROIs | Linear C=1 | MAE ± SD | 10.22 ± 7.05* | | 10.40 ± 7.41* | | 9.67 ± 6.89* | | 9.95 ± 6.83 | 10.32 ± 7.25 | 9.53 ± 6.66 |
|  |  | Perm p-value (NKI/HCPA) | 0.00 | 0.00 | 0.02 | 0.00 | 0.00 | 0.00 | - | - | - |
|  |  | Corr | 0.62 | | 0.59 | | 0.66 | | 0.64 | 0.61 | 0.68 |
|  | Poly2 C=1 | MAE ± SD | 10.41 ± 7.22 | | 10.52 ± 7.61 | | 9.82 ± 7.11 | | 10.38 ± 7.26 | 10.52 ± 7.41 | 9.84 ± 7.01 |
|  |  | Perm p-value (NKI/HCPA) | 0.02 | 0.52 | 0.09 | 0.00 | 0.19 | 0.61 | - | - | - |
|  |  | Corr | 0.58 | | 0.56 | | 0.63 | | 0.58 | 0.57 | 0.63 |
|  | RBF C=10000 | MAE ± SD | 9.87 ± 6.86* | | 10.23 ± 7.30* | | 9.65 ± 6.95 | | 9.74 ± 6.63 | 10.08 ± 7.21 | 9.50 ± 6.80 |
|  |  | Perm p-value (NKI/HCPA) | 0.00 | 0.00 | 0.01 | 0.00 | 0.01 | 0.09 | - | - | - |
|  |  | Corr | 0.64 | | 0.60 | | 0.65 | | 0.66 | 0.61 | 0.66 |
| CT+SC 425 ROIs | Linear C=1 | MAE ± SD | 10.17 ± 7.02* | | 10.32 ± 7.40* | | 9.55 ± 6.87* | | 9.88 ± 6.89 | 10.26 ± 7.24 | 9.45 ± 6.66 |
|  |  | Perm p-value (NKI/HCPA) | 0.00 | 0.00 | 0.00 | 0.00 | 0.00 | 0.00 | - | - | - |
|  |  | Corr | 0.62 | | 0.60 | | 0.66 | | 0.64 | 0.61 | 0.68 |
|  | Poly2 C=1 | MAE ± SD | 10.42 ± 7.21 | | 10.46 ± 7.56* | | 9.68 ± 7.06 | | 10.32 ± 7.23 | 10.49 ± 7.40 | 9.73 ± 6.97 |
|  |  | Perm p-value (NKI/HCPA) | 0.01 | 0.67 | 0.03 | 0.02 | 0.22 | 0.48 | - | - | - |
|  |  | Corr | 0.59 | | 0.56 | | 0.64 | | 0.59 | 0.57 | 0.64 |
|  | RBF C=10000 | MAE ± SD | 9.81 ± 6.82* | | 10.07 ± 7.18* | | 9.52 ± 6.92 | | 9.69 ± 6.60 | 9.90 ± 7.12 | 9.34 ± 6.87 |
|  |  | Perm p-value (NKI/HCPA) | 0.00 | 0.00 | 0.01 | 0.00 | 0.00 | 0.10 | - | - | - |
|  |  | Corr | 0.65 | | 0.62 | | 0.66 | | 0.66 | 0.63 | 0.67 |
| CT+SC+WM  497 ROIs | Linear C=1 | MAE ± SD | 10.01 ± 6.85* | | 10.17 ± 7.21* | | 9.38 ± 6.66*† | | 9.83 ± 6.74 | 10.12 ± 7.01 | 9.26 ± 6.56 |
|  |  | Perm p-value (NKI/HCPA) | 0.00 | 0.00 | 0.01 | 0.00 | 0.00 | 0.00 | - | - | - |
|  |  | Corr | 0.64 | | 0.62 | | 0.68 | | 0.65 | 0.63 | 0.69 |
|  | Poly2 C=1 | MAE ± SD | 10.29 ± 7.14 | | 10.37 ± 7.35* | | 9.63 ± 6.85 | | 10.25 ± 7.11 | 10.44 ± 7.16 | 9.57 ± 6.79 |
|  |  | Perm p-value (NKI/HCPA) | 0.00 | 0.63 | 0.03 | 0.01 | 0.29 | 0.49 | - | - | - |
|  |  | Corr | 0.60 | | 0.58 | | 0.65 | | 0.60 | 0.59 | 0.66 |
|  | RBF C=10000 | MAE ± SD | 9.67 ± 6.77*† | | 9.94 ± 6.94*† | | 9.43 ± 6.85* | | 9.60 ± 6.54† | 9.71 ± 6.92† | 9.22 ± 6.75† |
|  |  | Perm p-value (NKI/HCPA) | 0.00 | 0.00 | 0.01 | 0.00 | 0.02 | 0.01 | - | - | - |
|  |  | Corr | 0.66 | | 0.64 | | 0.67 | | 0.67 | 0.65 | 0.68 |

**Table S2.** Age prediction performance of the low-frequency fMRI physiological component variability (respiratory variation (RV), heart rate (HR), or RV and HR jointly) across linear, 2nd degree polynomial (Poly2) and radial basis function (RBF) kernel across different subsets of brain regions of interest (cortical - CT, subcortical - SC and white matter - WM ROIs). Age prediction accuracy is measured by mean absolute error (MAE, in years) and Pearson correlation coefficient (Corr) between chronological age and predicted age over two datasets.

* indicates when trained separately, the results achieved significant age prediction compared to the permutation test predicting age of the mismatched fMRI data (Perm p-value; p < 0.05) for both datasets. None of the permutation test results for predicting the age of mismatched physiological signals outperformed the original test (p = 0.00 for all), thus not shown in the table.

† marks the best age prediction performance (lowest MAE) for each column.

MAE ± SD, mean absolute error ± standard deviation of absolute error (in years).
