## Supplemental Table 3 for "Distributed fMRI patterns coupled to low-frequency cardiorespiratory dynamics provide markers of aging"

| NKI-MNI | Kernel | Metric | RV | | HR | | RVHR | | RAW-SD | RAW-fALFF | RAW-ALFF | REG-fALFF | REG-ALFF |
| --- | --- | --- | --- | --- | --- | --- | --- | --- | --- | --- | --- | --- | --- |
| CT 400 ROIs | Linear | MAE ± SD | 12.75 ± 8.50* | | 13.50 ± 8.94* | | 12.51 ± 8.62* | | 13.39 ± 11.65 | 12.53 ± 8.51 | 13.68 ± 11.19 | 12.17 ± 8.56 | 14.37 ± 12.71 |
|  | C=1 | Perm p-value | 0.00 | | 0.02 | | 0.00 | | 0.52 | 0.46 | 0.52 | 0.50 | 0.48 |
|  | Poly2 | MAE ± SD | 12.33 ± 8.30* | | 13.33 ± 9.13 | | 12.25 ± 8.47 | | 11.24 ± 8.01 | 9.83 ± 7.53† | 11.45 ± 8.00 | 9.75 ± 6.99† | 11.37 ± 8.24 |
|  | C=1 | Perm p-value | 0.02 | | 0.09 | | 0.19 | | 0.59 | 0.69 | 0.58 | 0.71 | 0.58 |
|  | RBF | MAE ± SD | 12.25 ± 8.32*† | | 13.15 ± 8.62*† | | 12.20 ± 8.29*† | | 10.47 ± 7.57† | 10.11 ± 6.84 | 10.43 ± 7.49† | 10.39 ± 6.97 | 10.44 ± 7.48† |
|  | C=10000 | Perm p-value | 0.00 | | 0.01 | | 0.01 | | 0.65 | 0.71 | 0.66 | 0.68 | 0.66 |
| NKI-IND | Kernel | Metric | RV | | HR | | RVHR | | RAW-SD | RAW-fALFF | RAW-ALFF | REG-fALFF | REG-ALFF |
| CT 400 ROIs | Linear | MAE ± SD | 13.20 ± 8.90* | | 13.96 ± 8.88 | | 13.05 ± 8.80 | | 15.13 ± 12.89 | 12.97 ± 8.82 | 15.25 ± 12.90 | 12.38 ± 8.71 | 17.04 ± 14.00 |
|  | C=1 | Perm p-value | 0.00 | | 0.26 | | 0.06 | | 0.37 | 0.39 | 0.39 | 0.47 | 0.32 |
|  | Poly2 | MAE ± SD | 13.11 ± 8.96 | | 14.05 ± 9.06 | | 13.39 ± 8.87 | | 11.91 ± 7.90 | 11.57 ± 8.85 | 12.01 ± 8.02 | 11.51 ± 8.25 | 11.94 ± 8.50 |
|  | C=1 | Perm p-value | 0.20 | | 0.54 | | 0.59 | | 0.55 | 0.53 | 0.54 | 0.57 | 0.54 |
|  | RBF | MAE ± SD | 12.80 ± 8.31* | | 13.74 ± 8.37 | | 12.75 ± 8.30* | | 11.30 ± 7.82 | 11.92 ± 7.85 | 11.65 ± 7.95 | 11.17 ± 7.36 | 11.60 ± 8.10 |
|  | C=10000 | Perm p-value | 0.01 | | 0.17 | | 0.04 | | 0.59 | 0.55 | 0.56 | 0.62 | 0.56 |

**Table S3.** Age prediction performance comparison between the common space MNI152-based cortical ROI extraction method (MNI) and the individual T1 structure-based ROI segmentation method (IND) across kernels (linear, 2nd degree polynomial - Poly2, and radial basis function - RBF).

* indicates when trained separately, the results achieved significant age prediction compared to the permutation test predicting age of the mismatched fMRI data (Perm p-value; p < 0.05) for both datasets. None of the permutation test results for predicting the age of mismatched physiological signals outperformed the empirical result (p = 0.00 for all), thus not shown in the table.

† marks the best age prediction performance (lowest MAE) for each column.

MAE ± SD, mean absolute error ± standard deviation of absolute error (in years); CT, cortical ROIs; RV, respiratory variation coupled fMRI variability; HR, heart rate coupled fMRI variability; RVHR, RV and HR jointly coupled fMRI variability; MAE ± SD, mean absolute error ± standard deviation of absolute error (in years); Corr, Pearson correlation coefficient.
