## Supplemental Table 4 for "Distributed fMRI patterns coupled to low-frequency cardiorespiratory dynamics provide markers of aging"

| NKI-MNI | Kernel | Metric | HR |  | Mixed-MNI | Kernel | Metric | HR |
| --- | --- | --- | --- | --- | --- | --- | --- | --- |
| CT 400 ROIs | RBF | MAE | 19.17* |  | CT 400 ROIs | RBF | MAE | 19.91 |
|  | C=10000 | *t*-test p-value | 0.0004 |  |  | C=10000 | *t*-test p-value | 0.0129 |
| CT+SC 425 ROIs | RBF | MAE | 18.80* |  | CT+SC 425 ROIs | RBF | MAE | 19.42* |
|  | C=10000 | *t*-test p-value | 0.0001 |  |  | C=10000 | *t*-test p-value | 0.0028 |
| CT+SC+WM 497 ROIs | RBF | MAE | 18.78* |  | CT+SC+WM 497 ROIs | RBF | MAE | 19.57 |
|  | C=10000 | *t*-test p-value | 0.0003 |  |  | C=10000 | *t*-test p-value | 0.0051 |

**Table S4.** Generalizability of the HR-fMRI variability age prediction performance trained on NKI (left), or NKI and HCPA combined (Mixed; right), and tested on HRV-ER dataset with radial basis function (RBF) kernel with different subsets of brain regions of interest (cortical - CT, subcortical - SC and white matter - WM ROIs). Prediction significance was determined with a one-tailed unpaired group-level *t*-test comparing the age prediction output between the older and young adult groups (* indicates p < 0.005). HR, heart rate coupled fMRI variability.
