## Supplemental Table 5 for "Distributed fMRI patterns coupled to low-frequency cardiorespiratory dynamics provide markers of aging"

| Component | Network Name | | VisCent | VisPeri | SomMotA | SomMotB | DorsAttnA | DorsAttnB | SalVentAttnA | SalVentAttnB | LimbicB | LimbicA | ContA | ContB | ContC | DefaultA | DefaultB | DefaultC | TempPar |
| --- | --- | --- | --- | --- | --- | --- | --- | --- | --- | --- | --- | --- | --- | --- | --- | --- | --- | --- | --- |
| RV | Linear C=1 | MAE | 12.60 | 12.25 | 11.81 | 11.98 | 12.31 | 12.20 | 11.46† | 12.37 | 12.51 | 12.40 | 12.00 | 12.39 | 12.53 | 11.79 | 11.88 | 12.37 | 12.50 |
|  |  | MAE SD | 8.57 | 8.25 | 8.22 | 8.09 | 8.13 | 8.24 | 7.70 | 8.18 | 8.22 | 8.27 | 8.06 | 8.17 | 8.54 | 7.84 | 7.84 | 8.31 | 8.54 |
|  |  | Corr | 0.22 | 0.34 | 0.40 | 0.38 | 0.34 | 0.34 | 0.49 | 0.33 | 0.31 | 0.34 | 0.40 | 0.32 | 0.26 | 0.46 | 0.44 | 0.32 | 0.24 |
|  | Poly2 C=1 | MAE | 12.48 | 12.27 | 11.87 | 11.95 | 12.32 | 12.17 | 11.37† | 12.24 | 12.54 | 12.25 | 11.86 | 12.49 | 12.31 | 11.69 | 11.71 | 12.30 | 12.53 |
|  |  | MAE SD | 8.48 | 8.17 | 8.19 | 8.10 | 8.13 | 8.27 | 7.72 | 8.02 | 8.39 | 8.30 | 8.02 | 8.18 | 8.49 | 7.85 | 7.98 | 8.35 | 8.53 |
|  |  | Corr | 0.25 | 0.32 | 0.37 | 0.38 | 0.32 | 0.33 | 0.48 | 0.35 | 0.26 | 0.32 | 0.41 | 0.29 | 0.29 | 0.43 | 0.42 | 0.30 | 0.23 |
|  | RBF C=1 | MAE | 12.25 | 12.18 | 11.92 | 11.83 | 12.25 | 12.21 | 11.46† | 12.20 | 12.44 | 12.46 | 11.94 | 12.35 | 12.35 | 11.82 | 11.91 | 12.34 | 12.34 |
|  |  | MAE SD | 8.47 | 8.13 | 8.23 | 7.99 | 7.99 | 8.19 | 7.70 | 8.09 | 8.33 | 8.27 | 8.08 | 8.24 | 8.46 | 7.91 | 7.89 | 8.27 | 8.34 |
|  |  | Corr | 0.34 | 0.37 | 0.39 | 0.43 | 0.38 | 0.35 | 0.51 | 0.37 | 0.29 | 0.31 | 0.42 | 0.32 | 0.31 | 0.46 | 0.44 | 0.33 | 0.30 |
| HR | Linear C=1 | MAE | 12.58 | 12.33 | 12.10 | 11.96† | 12.42 | 12.42 | 12.02 | 12.48 | 12.49 | 12.49 | 12.40 | 12.42 | 12.59 | 11.99 | 12.00 | 12.46 | 12.57 |
|  |  | MAE SD | 8.60 | 8.21 | 8.30 | 8.19 | 8.39 | 8.35 | 8.01 | 8.39 | 8.33 | 8.34 | 8.40 | 8.27 | 8.59 | 8.09 | 8.13 | 8.37 | 8.57 |
|  |  | Corr | 0.22 | 0.35 | 0.36 | 0.38 | 0.30 | 0.31 | 0.42 | 0.30 | 0.30 | 0.32 | 0.30 | 0.30 | 0.24 | 0.41 | 0.40 | 0.30 | 0.22 |
|  | Poly2 C=1 | MAE | 12.40 | 12.29 | 12.25 | 11.98 | 12.50 | 12.29 | 11.88† | 12.41 | 12.50 | 12.54 | 12.28 | 12.57 | 12.45 | 12.00 | 11.98 | 12.49 | 12.61 |
|  |  | MAE SD | 8.51 | 8.12 | 8.32 | 8.34 | 8.45 | 8.27 | 7.86 | 8.27 | 8.57 | 8.63 | 8.36 | 8.32 | 8.54 | 8.18 | 8.26 | 8.36 | 8.65 |
|  |  | Corr | 0.27 | 0.33 | 0.31 | 0.35 | 0.25 | 0.31 | 0.41 | 0.29 | 0.24 | 0.23 | 0.30 | 0.25 | 0.25 | 0.36 | 0.36 | 0.27 | 0.19 |
|  | RBF C=1 | MAE | 12.29 | 12.22 | 11.97 | 11.74† | 12.10 | 12.13 | 11.92 | 12.33 | 12.40 | 12.49 | 12.22 | 12.22 | 12.46 | 11.89 | 11.83 | 12.28 | 12.26 |
|  |  | MAE SD | 8.53 | 8.19 | 8.14 | 8.06 | 8.24 | 8.22 | 8.07 | 8.23 | 8.37 | 8.37 | 8.38 | 8.17 | 8.56 | 8.12 | 8.02 | 8.30 | 8.42 |
|  |  | Corr | 0.31 | 0.37 | 0.41 | 0.43 | 0.38 | 0.37 | 0.43 | 0.34 | 0.31 | 0.29 | 0.33 | 0.36 | 0.26 | 0.42 | 0.45 | 0.35 | 0.31 |
| RVHR | Linear C=1 | MAE | 12.45 | 12.14 | 11.60 | 11.78 | 12.18 | 12.05 | 11.46† | 12.19 | 12.42 | 12.28 | 11.90 | 12.23 | 12.44 | 11.61 | 11.58 | 12.23 | 12.39 |
|  |  | MAE SD | 8.37 | 8.08 | 8.06 | 8.02 | 7.93 | 7.94 | 7.51 | 7.92 | 8.10 | 8.13 | 7.94 | 7.94 | 8.44 | 7.69 | 7.65 | 8.14 | 8.37 |
|  |  | Corr | 0.28 | 0.37 | 0.44 | 0.41 | 0.38 | 0.39 | 0.50 | 0.39 | 0.33 | 0.37 | 0.42 | 0.36 | 0.30 | 0.48 | 0.47 | 0.37 | 0.29 |
|  | Poly2 C=1 | MAE | 12.24 | 12.09 | 11.64 | 11.81 | 12.18 | 11.98 | 11.29† | 11.99 | 12.48 | 12.21 | 11.72 | 12.36 | 12.25 | 11.43 | 11.45 | 12.13 | 12.41 |
|  |  | MAE SD | 8.31 | 7.92 | 8.05 | 8.12 | 8.01 | 7.86 | 7.44 | 7.67 | 8.28 | 8.28 | 8.02 | 7.98 | 8.31 | 7.73 | 7.76 | 8.14 | 8.37 |
|  |  | Corr | 0.32 | 0.38 | 0.42 | 0.39 | 0.35 | 0.39 | 0.50 | 0.41 | 0.29 | 0.33 | 0.41 | 0.34 | 0.32 | 0.47 | 0.46 | 0.36 | 0.28 |
|  | RBF C=1 | MAE | 12.02 | 12.12 | 11.62 | 11.52 | 11.96 | 11.97 | 11.33† | 11.93 | 12.25 | 12.32 | 11.76 | 12.13 | 12.31 | 11.48 | 11.44 | 12.12 | 12.10 |
|  |  | MAE SD | 8.23 | 8.02 | 8.00 | 7.86 | 7.78 | 7.89 | 7.54 | 7.90 | 8.26 | 8.21 | 7.98 | 8.03 | 8.41 | 7.81 | 7.65 | 8.07 | 8.17 |
|  |  | Corr | 0.39 | 0.38 | 0.44 | 0.46 | 0.43 | 0.41 | 0.52 | 0.42 | 0.33 | 0.34 | 0.44 | 0.37 | 0.31 | 0.49 | 0.49 | 0.38 | 0.36 |

**Table S5.** Schaefer atlas 17 network-level age prediction results across kernels (linear, 2nd degree polynomial - Poly2 and radial basis function - RBF).

† marks the best age prediction performance (lowest MAE) for each column for each fMRI physiological component.

RV, respiratory variation coupled fMRI variability; HR, heart rate coupled fMRI variability; RVHR, RV and HR jointly coupled fMRI variability; MAE SD, standard deviation of mean absolute error (in years); Corr, Pearson correlation coefficient. VisCent - central visual network; SomMot - somatomotor network; DorsAttn - dorsal attention network; SalVentAttn - salience / ventral attention network; Limbic - limbic network; Cont - control network; Default - default mode network; TempPar - temporal parietal network.
