## Supplemental Table 6 for "Distributed fMRI patterns coupled to low-frequency cardiorespiratory dynamics provide markers of aging"

| NKI+HCPA-MNI | Kernel | Metric | RAW-SD | RAW-fALFF | RAW-ALFF | REG-fALFF | REG-ALFF | RAW-SD (Joint) | RAW-fALFF  (Joint) | RAW-ALFF (Joint) | REG-fALFF  (Joint) | REG-ALFF  (Joint) |
| --- | --- | --- | --- | --- | --- | --- | --- | --- | --- | --- | --- | --- |
| CT  400 ROIs | Linear | MAE ± SD | 10.84 ± 8.66 | 9.04 ± 6.66 | 11.96 ± 10.52 | 9.08 ± 6.65 | 11.27 ± 9.19 | 15.53 ± 15.07 | 9.03 ± 6.69 | 236.31 ± 267.52 | 9.18 ± 6.74 | 219.11 ± 236.12 |
|  | C=1 | Corr | 0.61 | 0.70 | 0.55 | 0.70 | 0.57 | 0.21 | 0.72 | 0.06 | 0.71 | 0.08 |
|  | Poly2 | MAE ± SD | 8.88 ± 6.81 | 8.34 ± 6.24 | 9.21 ± 7.06 | 8.37 ± 6.16 | 8.99 ± 6.89 | 10.18 ± 8.30 | 8.61 ± 6.49 | 10.44 ± 8.79 | 8.73 ± 6.47 | 10.20 ± 8.57 |
|  | C=1 | Corr | 0.70 | 0.74 | 0.67 | 0.74 | 0.69 | 0.55 | 0.72 | 0.50 | 0.72 | 0.53 |
|  | RBF | MAE ± SD | 8.47 ± 6.22 | 8.41 ± 6.11 | 8.39 ± 6.24 | 8.52 ± 6.17 | 8.38 ± 6.05 | 8.66 ± 6.83 | 8.38 ± 6.10 | 8.40 ± 6.40 | 8.53 ± 6.27 | 8.31 ± 6.20 |
|  | C=10000 | Corr | 0.74 | 0.75 | 0.75 | 0.74 | 0.75 | 0.70 | 0.75 | 0.74 | 0.73 | 0.75 |
| CT+SC 425 ROIs | Linear | MAE ± SD | 10.31 ± 8.38 | 8.92 ± 6.61 | 11.76 ± 9.61 | 9.01 ± 6.60 | 10.91 ± 8.45 | 14.00 ± 13.32 | 8.81 ± 6.49 | 159.72 ± 130.11 | 9.00 ± 6.57 | 163.16 ± 150.67 |
|  | C=1 | Corr | 0.64 | 0.71 | 0.57 | 0.70 | 0.59 | 0.28 | 0.73 | 0.08 | 0.72 | 0.08 |
|  | Poly2 | MAE ± SD | 8.42 ± 6.43 | 8.09 ± 6.03 | 8.81 ± 6.61 | 8.17 ± 5.99 | 8.56 ± 6.47 | 9.80 ± 8.17 | 8.46 ± 6.40 | 10.04 ± 8.48 | 8.51 ± 6.40 | 9.84 ± 8.35 |
|  | C=1 | Corr | 0.73 | 0.76 | 0.71 | 0.76 | 0.72 | 0.59 | 0.73 | 0.55 | 0.73 | 0.57 |
|  | RBF | MAE ± SD | 7.95 ± 6.10 | 8.24 ± 6.01 | 8.01 ± 6.05 | 8.32 ± 6.04 | 7.93 ± 5.89 | 8.12 ± 6.51 | 8.22 ± 5.97 | 8.05 ± 6.13 | 8.33 ± 6.10 | 7.91 ± 6.00 |
|  | C=10000 | Corr | 0.77 | 0.76 | 0.77 | 0.75 | 0.78 | 0.74 | 0.76 | 0.76 | 0.75 | 0.77 |
| CT+SC+WM 497 ROIs | Linear | MAE ± SD | 9.97 ± 8.13 | 8.73 ± 6.45 | 11.10 ± 8.87 | 8.90 ± 6.52 | 10.65 ± 8.46 | 13.25 ± 12.83 | 8.67 ± 6.36 | 180.73 ± 170.69 | 8.85 ± 6.45 | 175.85 ± 176.72 |
|  | C=1 | Corr | 0.66 | 0.72 | 0.58 | 0.71 | 0.62 | 0.33 | 0.75 | 0.08 | 0.74 | 0.03 |
|  | Poly2 | MAE ± SD | 8.35 ± 6.33 | 7.89 ± 5.85† | 8.79 ± 6.49 | 8.09 ± 5.83† | 8.60 ± 6.40 | 9.76 ± 8.22 | 8.25 ± 6.24 | 10.08 ± 8.49 | 8.39 ± 6.32 | 9.90 ± 8.41 |
|  | C=1 | Corr | 0.74 | 0.77 | 0.71 | 0.77 | 0.72 | 0.59 | 0.75 | 0.54 | 0.74 | 0.57 |
|  | RBF | MAE ± SD | 7.84 ± 6.05† | 8.13 ± 5.90 | 7.87 ± 6.03† | 8.15 ± 5.81 | 7.79 ± 5.86† | 8.00 ± 6.37† | 8.13 ± 5.88† | 8.01 ± 6.06† | 8.17 ± 5.86† | 7.88 ± 5.95† |
|  | C=10000 | Corr | 0.77 | 0.77 | 0.78 | 0.77 | 0.78 | 0.75 | 0.77 | 0.77 | 0.77 | 0.77 |

**Table S6.** Age prediction performance of the non-physiological fMRI metrics (original fMRI data - RAW, fMRI data with low-frequency respiratory variation and heart rate effects regressed out - REG,  standard deviation - SD, amplitude of low frequency fluctuations - ALFF, fractional amplitude of low frequency fluctuations - fALFF) across linear, 2nd degree polynomial (Poly2) and radial basis function (RBF) kernel with different subsets of brain regions of interest (cortical - CT, subcortical - SC and white matter - WM ROIs).

† marks the best age prediction performance (lowest mean absolute error; MAE, in years) for each column.

MAE ± SD, mean absolute error ± standard deviation of absolute error (in years); Corr, Pearson correlation coefficient.
