## Supplemental Table 7 for "Distributed fMRI patterns coupled to low-frequency cardiorespiratory dynamics provide markers of aging"

| NKI+HCPA-MNI | Kernel | Metric | RV | HR | RVHR | RV(Joint) | HR(Joint) | RVHR(Joint) |
| --- | --- | --- | --- | --- | --- | --- | --- | --- |
| CT+SC+WM   497 ROIs +  mean FD | Linear C=1 | MAE ± SD | 10.02 ± 6.82 | 10.17 ± 7.18 | 9.37 ± 6.63† | 9.82 ± 6.73 | 10.13 ± 7.00 | 9.26 ± 6.56 |
|  |  | *t*-test t-value | 0.49 | -0.89 | -2.11* | -0.13 | 0.78 | 0.39 |
|  |  | *t*-test p-value | 0.62 | 0.37 | 0.04 | 0.89 | 0.44 | 0.70 |
|  |  | Corr | 0.64 | 0.62 | 0.68 | 0.65 | 0.63 | 0.69 |
|  | Poly2 C=1 | MAE ± SD | 10.29 ± 7.12 | 10.36 ± 7.32 | 9.58 ± 6.78 | 10.23 ± 7.09 | 10.44 ± 7.13 | 9.56 ± 6.77 |
|  |  | *t*-test t-value | -0.67 | -1.02 | -3.09* | -1.24 | 0.61 | -1.11 |
|  |  | *t*-test p-value | 0.51 | 0.31 | <0.01 | 0.22 | 0.54 | 0.27 |
|  |  | Corr | 0.60 | 0.59 | 0.66 | 0.60 | 0.59 | 0.66 |
|  | RBF C=10000 | MAE ± SD | 9.65 ± 6.72† | 9.92 ± 6.89† | 9.39 ± 6.80 | 9.60 ± 6.50† | 9.73 ± 6.90† | 9.20 ± 6.72† |
|  |  | *t*-test t-value | -1.70 | -1.00 | -2.69* | -0.20 | 1.06 | -1.77 |
|  |  | *t*-test p-value | 0.09 | 0.32 | 0.01 | 0.84 | 0.29 | 0.08 |
|  |  | Corr | 0.66 | 0.64 | 0.67 | 0.68 | 0.65 | 0.69 |

| NKI+HCPA-MNI | Kernel | Metric | RAW | RAW-fALFF | RAW-ALFF | REG-fALFF | REG-ALFF | RAW(Joint) | RAW-fALFF (Joint) | RAW-ALFF  (Joint) | REG-fALFF  (Joint) | REG-ALFF  (Joint) |
| --- | --- | --- | --- | --- | --- | --- | --- | --- | --- | --- | --- | --- |
| CT+SC+WM   497 ROIs + mean FD | Linear C=1 | MAE ± SD | 9.98 ± 8.03 | 8.69 ± 6.41 | 11.10 ± 8.87 | 8.87 ± 6.49 | 10.71 ± 8.51 | 14.65 ± 15.06 | 8.66 ± 6.37 | 188.50 ± 199.88 | 8.85 ± 6.46 | 214.90 ± 204.70 |
|  |  | *t*-test t-value | 0.05 | -2.05* | 1.11 | -2.17* | 0.38 | 5.17* | -0.65 | 1.56 | -0.15 | 6.41* |
|  |  | *t*-test p-value | 0.96 | 0.04 | 0.27 | 0.03 | 0.70 | <0.01 | 0.52 | 0.12 | 0.88 | <0.01 |
|  |  | Corr | 0.66 | 0.73 | 0.58 | 0.72 | 0.62 | 0.25 | 0.75 | 0.10 | 0.74 | 0.04 |
|  | Poly2 C=1 | MAE ± SD | 8.35 ± 6.33 | 7.84 ± 5.73† | 8.79 ± 6.49 | 7.96 ± 5.81† | 8.60 ± 6.40 | 9.76 ± 8.22 | 8.24 ± 6.23 | 10.09 ± 8.49 | 8.40 ± 6.35 | 9.90 ± 8.41 |
|  |  | *t*-test t-value | 4.98* | -1.32 | 2.34* | -2.93* | 1.77 | 6.13* | -0.23 | 2.50* | 0.17 | 3.57* |
|  |  | *t*-test p-value | <0.01 | 0.19 | 0.02 | <0.01 | 0.08 | <0.01 | 0.82 | 0.01 | 0.86 | <0.01 |
|  |  | Corr | 0.74 | 0.78 | 0.71 | 0.77 | 0.72 | 0.59 | 0.75 | 0.54 | 0.74 | 0.57 |
|  | RBF C=10000 | MAE ± SD | 7.84 ± 6.05† | 8.12 ± 5.89 | 7.87 ± 6.03† | 8.10 ± 5.80 | 7.79 ± 5.86† | 8.00 ± 6.37† | 8.15 ± 5.87† | 8.01 ± 6.06† | 8.13 ± 5.87† | 7.88 ± 5.96† |
|  |  | *t*-test t-value | -0.62 | -0.24 | -2.32* | -1.85 | -2.17* | 0.99 | 0.58 | 4.07* | -1.26 | 3.11* |
|  |  | *t*-test p-value | 0.54 | 0.81 | 0.02 | 0.06 | 0.03 | 0.32 | 0.56 | <0.01 | 0.21 | <0.01 |
|  |  | Corr | 0.77 | 0.77 | 0.78 | 0.77 | 0.78 | 0.75 | 0.77 | 0.76 | 0.77 | 0.77 |

**Table S7.** Effect of including head motion (mean framewise displacement; mean FD) as an additional feature in age-prediction models. Age-prediction performance was assessed across different SVR kernels (linear, 2nd degree polynomial (Poly2) and radial basis function (RBF)), fMRI physiological components (respiratory variation (RV), heart rate (HR), or RV and HR together (RVHR)), non-physiological fMRI metrics (original fMRI data - RAW, fMRI data with low-frequency respiratory variation and heart rate effects regressed out - REG,  standard deviation - SD, amplitude of low frequency fluctuations - ALFF, fractional amplitude of low frequency fluctuations - fALFF) and training strategies. Statistical significance was assessed using a two-tailed paired *t*-test on age prediction MAE, averaged across segments within a given subject, between the model including mean FD and the model without mean FD (bottom row of **Table S1 & S2**); positive t-values indicate higher MAE (i.e., worse age prediction performance) for the model including mean FD relative to the model without mean FD.

* indicates significant differences (p < 0.05).

† marks the best age prediction performance (lowest mean absolute error; MAE, in years) for each column.

Corr, Pearson correlation coefficient; Joint, training jointly on both NKI and HCPA datasets.
