## Supplemental Table 8 for "Distributed fMRI patterns coupled to low-frequency cardiorespiratory dynamics provide markers of aging"

Measures

NIH Toolbox Fluid Cognition Composite Score (Age-Adjusted)

NIH Toolbox Crystallized Cognition Composite Score (Age-Adjusted)

Penn Emotion Recognition Test (ER40) – Total Correct Responses

NIH Toolbox Flanker Inhibitory Control and Attention Test (Age-Adjusted Score)

NIH Toolbox Ability to Participate in Social Roles and Activities (Fixed Form) Score

NIH Toolbox Oral Reading Recognition Test (Age-Adjusted Score)

NIH Toolbox Pattern Comparison Processing Speed Test (Age-Adjusted Score)

NIH Toolbox Negative Affect – Emotional Distress T-score

PROMIS Global Health T-score

NIH Toolbox Social Relationships – Social Isolation T-score

NIH Toolbox Picture Sequence Memory Test (Age-Adjusted Score)

NIH Toolbox Self-Efficacy (Computer Adaptive Test) T-score

NIH Toolbox Friendship (Fixed Form) T-score

NIH Toolbox Hostility (Fixed Form) T-score

NIH Toolbox Perceived Rejection (Fixed Form) T-score

Penn Temporal Pole Visual Perception (TPVP) – Age-Corrected Score

OASR Aggressive Behavior Scale

OASR Anxious/Depressed Scale

OASR Attention Problems Scale

OASR Rule-Breaking Behavior Scale

OASR Somatic Complaints Scale

OASR Thought Problems Scale

OASR Withdrawn Scale

OASR Intrusive Behavior Scale

**Table S8.** List of cognitive and behavioral metrics investigated in relation to the corrected brain age gap on the HCPA dataset. The corrected brain age gap was computed from the age prediction using physio-fMRI variability metrics (respiratory variation (RV), heart rate (HR), or RV and HR together (RVHR)), or fMRI variability with physiological components regressed out. OASR, older adult self report.
