## Supplemental Text 1 for "Distributed fMRI patterns coupled to low-frequency cardiorespiratory dynamics provide markers of aging"

**Text S1**

**Impact of model nonlinearities and regularization**

The effects of kernel and associated kernel nonlinearity on age prediction accuracy have not been extensively studied in the fMRI variability and aging community. In our study, we explored kernel effects by comparing 3 SVR kernels with different levels of nonlinearity (linear, second-degree polynomial, and RBF kernels). We observed that the nonlinear RBF kernel achieved the best age prediction performance with RV- or HR-fMRI variability, whereas the linear kernel obtained similar or better accuracy on the jointly fitted RVHR-fMRI variability compared to the RBF kernel. In related work using non-physiological imaging features, Schulz et al. have investigated the influence of model complexity and kernel nonlinearity on age/sex prediction using structural and functional connectivity measures, and found that in the case of limited (here <10,000) sample size, the linear kernel achieved “indistinguishable” performance compared to nonlinear kernels (Schulz et al. 2020). This observation suggests that in the face of noisy and limited data (which may be the case for the present study), a linear model can learn effectively even if the underlying relationship is nonlinear, potentially explaining the strong performance of the linear kernel that we observe in the case of RVHR-fMRI variability. A second possibility is that the relationship between the fMRI physiological component and age is truly linear (perhaps with a small nonlinear component). In another study using subject-level phenotypic measures across multiple organ systems to predict chronological age (including pulmonary and cardiovascular systems), it has been found that the RBF kernel did not improve age prediction accuracy compared to the linear kernel (Tian et al. 2023). If a linear relationship also dominates in the case of the dynamic physiological components captured in fMRI by RV and HR, then perhaps the joint effect of modeling RV and HR captures more completely a latent, underlying component of the fMRI signal linked with autonomic physiology (Bolt et al. 2025), leading to the superior performance of the linear model for this case; while for RV and HR individually, the relationship with age may be more complex and nonlinear. However, the current study only weakly supports this claim, and further testing is needed to confirm it. Future studies could test this hypothesis by including additional dynamic physiological signals (e.g. blood pressure or skin conductance time courses concurrently recorded with fMRI) into the age prediction framework and seeing if the linear kernel performance is more similar to, or surpasses, that of the RBF kernel.

With regard to model hyperparameters an epsilon tube of 0.01 and gamma (hyperparameter for nonlinear kernels only) set to ‘scale’ (1 / (number of features * overall feature variance), which equals about 0.1) obtained the best age prediction performance on the validation set. The main hyperparameter that differed across the three kernels used in the SVR model was the regularization term C, where the regularization strength is inversely proportional to C: in other words, the smaller the C, the stronger the L2 regularization. The linear and 2nd degree polynomial kernels were optimized with C=1, and the RBF kernel was optimized with C=10000. For the RBF kernel, the optimal C value is closely linked with the gamma parameter. A lower gamma value for the RBF kernel, which is the case for our model, usually serves as a good structural regularizer itself and produces a smoother function. By pairing a relatively small gamma value with less L2 regularization (larger C), the RBF-based SVR model can be made more complex thus capturing data complexity and preventing underfitting (PedregosaFabian et al. 2011). However, this effect is much less profound with the Poly2 kernel, where increasing gamma to potentially improve model complexity did not improve validation set performance.
