## Supplemental Text 2 for "Distributed fMRI patterns coupled to low-frequency cardiorespiratory dynamics provide markers of aging"

**Text S2**

**Potential role of age-related brain anatomy changes**

Brain morphology can change substantially across the adult lifespan. This could potentially lead to brain registration errors in the older adults, which might impact age prediction results with fMRI variability. To investigate this possibility, we compared two ROI extraction strategies in the NKI dataset: 1) registering individual structural images to a common MNI152 space and extracting ROIs from this common space, and 2) extracting subject-specific ROIs in native space using FreeSurfer’s surface-based segmentation. We found that the MNI152 common-space approach outperformed the individualized approach in predicting age – not only from the component of fMRI data linked with physiological variability, but also for BOLD variability calculated from the raw and regressed signals. One explanation is that by first registering the brain structures onto a common template, the ROIs were better aligned with one another. Yet, it is also possible that any systematic age-related differences in MNI registration accuracy might inflate the age prediction ability of fMRI data. However, the MNI-based results for fMRI-physiological variability were significant compared with our permutation tests, which theoretically control for potential brain registration error as well, suggesting that registration error may not be the primary reason for the higher age prediction performance achieved by the MNI-based approach.

On the other hand, though FreeSurfer’s native-space segmentation avoids volumetric warping related registration errors, it may still be affected by age-related differences in cortical geometry. Because the Schaefer atlas was originally derived from young adults, age-related changes in cortical folding patterns can reduce the accuracy when matching the atlas to older adults’ native anatomical surfaces. This reduced correspondence might contribute to the comparatively lower performance of native-space ROI features in predicting age.
