## Supplemental Text 3 for "Distributed fMRI patterns coupled to low-frequency cardiorespiratory dynamics provide markers of aging"

**Text S3**

**Age prediction performance with motion features**

While not our main goal, we tested the age prediction performance using the same 497 cortical, subcortical and white matter ROI variability values with an added motion feature (mean framewise displacement, or mean FD) (498 features in total; **Table S7**) with the exact same train/test/validation split and model hyperparameters. We did observe marginal age prediction performance improvement for some physio-fMRI variability models, and only the RVHR-fMRI variability model, when trained separately, showed significant improvement in age prediction accuracy after incorporating the motion feature (mean framewise displacement) (**Table S7**).
