## Supplemental Text 4 for "Distributed fMRI patterns coupled to low-frequency cardiorespiratory dynamics provide markers of aging"

**Text S4**

**Brain age estimated from fMRI variability metrics and cognitive or behavioral measures**

Grady et al. linked resting state fMRI signal standard deviation (SD) to cognitive and behavioral measures and found positive correlation between overall BOLD SD and positive socioemotional scores and morning chronotype, and negatives correlations with emotional scores and fluid cognition (Grady et al. 2023). On the other hand, brain age gap, or the difference between predicted age and chronological age, has been widely used to characterize abnormal or accelerated aging (Grady et al. 2023; Cumplido-Mayoral et al. 2023; Gaser et al. 2024; Goodman et al. 2024; Kim et al. 2025).

On top of our current age prediction framework, we followed the pipeline proposed in Gaser et al. to compute the corrected brain age gap for each subject in the HCPA dataset by 1) subtracting chronological age from predicted age (averaged across age predictions for each subject), and 2) fitting the linear age coefficients on the 10 validation sets and using the fitted coefficients to correct for brain age bias on the 10 corresponding test set separately. We then calculated Spearman rank correlation coefficient between corrected brain age gap with behavioral and cognitive metrics including 7 out of 8 older adult self-report (OASR) syndrome scales, and age-adjusted cognitive composite scores (listed in **Table S8**) for adults older than 60 years old (193 subjects).

Our preliminary exploration showed that none of the relationships survived FDR correction. The following are ones that have uncorrected p<0.05: OASR thought problems with RV-fMRI variability (r=0.191, p=0.008, FDR-corrected p=0.189), NIH Toolbox Ability to Participate in Social Roles and Activities (Fixed Form) Score with REG-fALFF (r=-0.150, p=0.037, FDR-corrected p=0.831), and REG-ALFF (r=-0.172, p=0.017, FDR-corrected p=0.409).
